# Ontogeny drives stage-specific effects of a Gata1 mutation

**DOI:** 10.1101/2025.01.15.633164

**Authors:** Alina Sommer, Stephan Fischer, Laina Freyer, Yvan Lallemand, Pascal Dardenne, Elisa Gomez Perdiguero

## Abstract

The biography of an organism, including, for instance, its genetic background and exposure to environmental influences, determines the penetrance and expressivity of monogenic diseases between individuals. This also applies at the single-cell level. Interestingly, mature blood and immune cells originate from anatomically and temporally distinct stem and progenitor populations at different stages of development. Here we show that the ontogeny or progenitor origin of megakaryocytes determines their developmental trajectory at the cellular and transcriptional level. As a result, a mutation in the key transcription factor Gata1 dysregulates megakaryopoiesis in an ontogeny-specific manner causing a stage-specific disorder. Particularly, only early extraembryonic hematopoietic progenitors drive blast-like cell development leading to a transient accumulation of megakaryocytes in the fetus but not in the adult. Therefore, pediatric blood disorders may differ functionally from adult diseases due to changing progenitor sources of mature blood and immune cells throughout development.

**Highlights:**

- Fetal and adult megakaryopoiesis pass through distinct immunophenotypic stages
- Ontogeny of megakaryocytes dictates their unique transcriptional trajectory
- Progenitor origin rather than the differentiation pathway determines the effect of mutant Gata1
- Block in maturation at the progenitor level causes the accumulation of yolk sac-derived megakaryocytes in the presence of mutant Gata1
- Mutant Gata1 drives blast cell formation exclusively in yolk sac-derived lineages

## Introduction

Depending on the genetic background (intrinsic) and environmental factors (extrinsic), the same mutation can exert different phenotypes between individuals (Nadeau, 2001; Cooper et al., 2013; Taeubner et al., 2018). This has been reported for instance for diseases such as breast cancer or Retinoblastoma (Onadim et al., 1992; Antoniou et al., 2003; King, Marks & Mandell, 2003). Similarly, within an organism, the intrinsic features and environmental signals a single cell experiences change during the development from embryo to adult which could explain the stage-specific effects of somatic mutations.

For example, blood and immune cells at different developmental stages trace back to distinct ontogenies and anatomical origins, as the hematopoietic system is formed from successive and overlapping progenitor waves. The first hematopoietic cells are produced from transient primitive cells and transient definitive erythro-myeloid progenitors (EMPs) that arise from the extraembryonic yolk sac (Palis et al., 1999; McGrath et al., 2015; Gomez Perdiguero et al., 2015; Sommer & Gomez Perdiguero, 2024). Around Embryonic day (E)14.5, hematopoiesis is then taken over by hematopoietic stem and progenitor cells (HSPCs) that emerge at midgestation in mice from intraembryonic hemogenic endothelium in the dorsal aorta (Müller et al., 1994; Medvinsky & Dzierzak, 1996; North et al., 1999; Iturri et al., 2021; Yokomizo et al., 2022). The molecular mechanisms underlying the emergence, commitment, and differentiation processes of EMPs and HSPCs differ between progenitor waves (Hadland et al., 2004; Chen et al., 2011; Schulz et al., 2012; Silvério-Alves et al., 2023). In particular, GATA1, a key transcription factor in megakaryopoiesis and erythropoiesis, has stage-specific functions and drives disorders specifically in fetal development when mutated (Shivdasani et al., 1997; Li et al., 2005). However, it is not yet fully understood to what extent extrinsic or intrinsic factors cause this stage-specific role of GATA1 (Miyauchi, 2024).

Using lineage tracing and single-cell multiome sequencing, we here show that EMP- and HSPC-derived megakaryopoiesis pass through distinct trajectories at the cellular level which are governed by different transcriptional programs. As a result, mutant Gata1 (*Gata1^mCherry^*) affects yolk sac lineages more severely than intraembryonic-derived HSPCs. Functionally, this translates into the accumulation of megakaryocytes derived from primitive cells and EMPs, but not HSPCs. In addition, exclusively yolk sac-derived cells give rise to CD244^+^ blast-like cells driven by the increased activity of the transcription factors CEBPB and Helios (*Ikzf2*) in *Gata1^mCherry^*compared to wild-type conditions.

Collectively, our results show that ontogeny, or the progenitor origin, is a key driver in the stage-specific function of GATA1. This will pave the way to investigate the functional consequences of ontogeny in disease development which may lead to a better understanding of the differences between pediatric and adult diseases.

## Results

### Fetal and adult megakaryopoiesis progress through immunophenotypically distinct progenitor stages

We first characterized the different stages of megakaryocyte differentiation and maturation throughout embryonic development and across different fetal hematopoietic niches, focusing on the extraembryonic yolk sac (at embryonic day (E) 10.5) and the fetal liver (from E12.5 to E16.5), the main hematopoietic niche in development. Adult bone marrow (BM) cells were used as a reference, where megakaryocyte progenitors (MkP) are defined as Lin^neg^ Kit^high^ Sca-1^neg^ CD41^+^ CD150^+^ cells (Pronk et al., 2007) and megakaryocytes (Mk) as CD41^high^ CD42^high^ Kit^neg^ (**Fig. 1A, Supplemental Data Fig. 1A**) (Coller et al., 1983). During fetal stages, no immunophenotypic MkPs were detected (**Supplemental Data Fig. 1A**).

**Figure 1.**
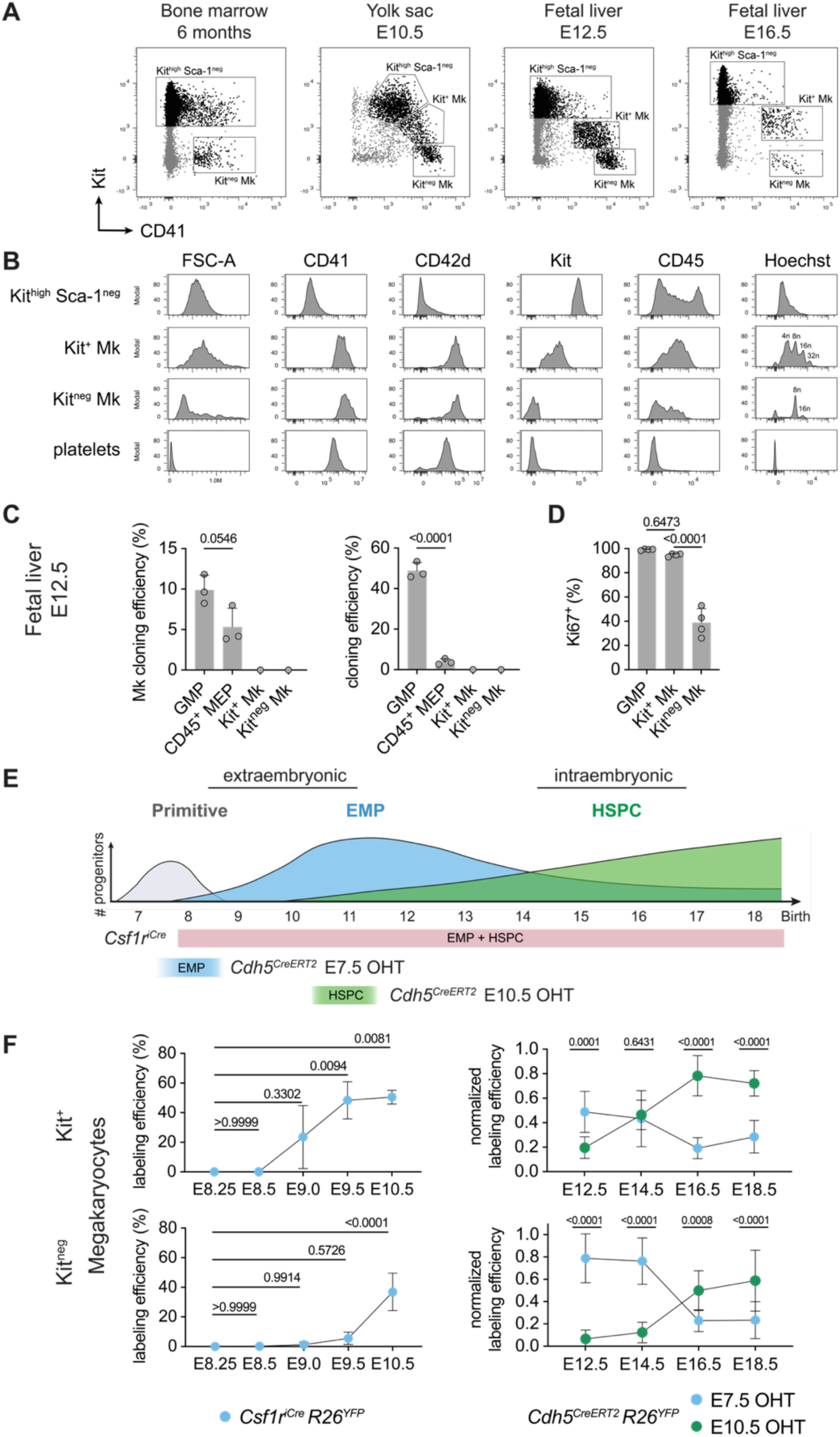
Stages and origins of megakaryopoiesis in fetal development. (**A**) Dot plots displaying Lin^neg^ Sca-1^neg^ cells from adult bone marrow (far left), E10.5 yolk sac (left), E12.5 (right), and E16.5 (far right) fetal liver. Three populations were defined based on the expression of Kit and CD41: Kit^high^ Sca-1^neg^ progenitors, Kit^+^ Mk, and Kit^neg^ Mk. (**B**) Histograms showing size (FSC-A), expression of CD41, CD42d, Kit, CD45, and ploidy (Hoechst 33342) of Kit^high^ Sca-1^neg^ progenitors, Kit^+^ and Kit^neg^ Mks and platelets in the E12.5 fetal liver as gated in **A**. (**C**) Bar plots showing the overall cloning efficiency (left) and Mk cloning efficiency (right) of immunophenotypic granulocyte-monocyte progenitors (GMPs, Lin^neg^ Kit^high^ Sca-1^neg^ CD16/32^high^ CD34^+^), CD45+ megakaryocyte-erythrocyte progenitors (MEPs, Lin^neg^ Kit^high^ Sca-1^neg^ CD16/32^neg^ CD34^neg^), Kit^+^ CD41^+^ Mks and Kit^neg^ CD41^+^ Mks sorted form E12.5 fetal livers and cultured for 7 days in liquid culture. 3 independent litters. Šidak multiple comparisons. (**D**) Bar plot showing the frequency of Ki67^+^ cells among GMPs, Kit^+^, and Kit^neg^ Mks in the E12.5 fetal liver. 4 embryos from one litter. Tukey’s multiple comparisons. (**E**) Scheme representing the contribution of the four main hematopoietic waves emerging in embryonic development (primitive cells, erythro-myeloid progenitors (EMP), and hematopoietic stem and progenitor cells (HSPC)) and strategies to fate-map them. *Csf1r^iCre^* labels all definitive hematopoietic progenitors and myeloid cells. *Cdh5^CreERT2^* pulsed with 4-OHT at E7.5 labels EMPs and their progeny whereas when pulsed at E10.5 HSPCs and their progeny are labeled. (**F**) Graph illustrating the labeling efficiency of Kit^+^ (top left) and Kit^neg^ (bottom left) Mks in E8.25-E10.5 yolk sacs of *Csf1r^iCre^ R26^YFP^* embryos (EMP contribution). Normalized labeling efficiency of Kit^+^ (top right) and Kit^neg^ (bottom right) Mks in E12.5-E18.5 fetal livers of *Cdh5^CreERT2^ R26^YF^*^P^ embryos pulsed with 4-OHT at E7.5 (blue, EMP contribution, normalized to microglia) or E10.5 (green, HSPC contribution, normalized to Lin^neg^ Sca-1^+^ Kit^high^). Left: One-way ANOVA with E8.25 as control, P-value = 0.0004 (top), <0.0001 (bottom). Dunnett’s multiple comparisons. Right: Two-way ANOVA. Embryonic stage: P-value <0.0001 (top), = 0.5063 (bottom) Ontogeny = 0.0087 (top), <0.0001 (bottom), P-value Interaction: P-value <0.0001 (top), <0.0001 (bottom). Tukey’s multiple comparisons. n ≥3 embryos from N ≥2 litters per condition. Data are represented as mean ± SD. See also Supplemental Data Fig. 1.

In contrast, two CD41^+^ populations could be distinguished in fetal hematopoietic tissues based on Kit and CD41 expression levels: Kit^+^ CD41^+^ and Kit^neg^ CD41^+^ (**Fig. 1A**). At E12.5, both Kit^+^ CD41^+^ and Kit^neg^ CD41^+^ cells expressed comparable levels of CD42d (platelet glycoprotein V) and CD41 (platelet glycoprotein IIb, *Itga2b*) (**Fig. 1B**). While Kit^+^ CD41^+^ cells displayed intermediate to low levels of CD45 and Kit, Kit^neg^ CD41^+^ cells were mostly negative for both markers. In regards to their ploidy, a unique feature of megakaryocytes, most Kit^+^ CD41^+^ cells were 4n, 8n, or 16n, whereas most Kit^neg^ cells were mainly 8n (**Fig. 1B**).

To investigate whether CD41^+^ cells were endowed with progenitor potential or rather corresponded to Mk maturation stages, we cultured single Kit^+^ and Kit^neg^ CD41^+^ cells in liquid and semi-solid medium. Neither Kit^+^ nor Kit^neg^ CD41^+^ cells were able to produce Mk colonies indicating that they lack clonogenic potential and are not progenitors (**Fig. 1C**). Hence, we named them Kit^+^ and Kit^neg^ Mk thereafter. Finally, virtually all Kit^+^ Mks were Ki67^+^ and at a similar frequency as granulocyte-monocyte progenitors (GMPs), whereas only 40% of Kit^neg^ Mks were Ki67^+^ (**Figure 1D**). The apparent contradiction between the inability to clone *in vitro* and the important proliferative capacity of Kit^+^ CD41^+^ cells can be accounted for by endomitosis. Endomitosis is a unique feature of megakaryopoiesis, where Mks increase their ploidy by undergoing several rounds of nuclear replication without cell division (Vitrat et al., 1998). Thus, Kit^+^ Mks corresponded to maturing Mks with increasing ploidy levels, whereas the smaller Kit^neg^ Mks corresponded to mature Mks.

To identify the progenitor upstream of Kit^+^ CD41^+^ cells endowed with Mk potential in the E12.5 fetal liver, we functionally compared GMPs (immunophenotypic equivalent to EMPs in the yolk sac, Lin^neg^ Kit^high^ Sca-1^neg^ CD34^+^ CD16/32^high^) and megakaryocyte-erythrocyte progenitors (MEP, Lin^neg^ Kit^high^ Sca-1^neg^ CD34^neg^ CD16/32^neg^) (**Fig. 1C**). Potency assays demonstrated that GMPs were the fetal hematopoietic population endowed with the strongest MkP potential (with regard to Mk cloning efficiency and number of Mks per colony), particularly when compared to CD45^+^ MEPs (**Fig. 1C & Supplemental Data Fig. 1B**).

Collectively, time course analysis showed fetal Mks (Kit^neg^ CD41^+^) are produced from Lin^neg^ Kit^high^ Sca-1^neg^ CD34^+^ CD16/32^high^ progenitors through an MEP stage only in the fetal liver. Fetal megakaryopoiesis differs from its adult counterpart due to the absence of Kit^high^ CD41^+^ CD150^+^ megakaryocyte progenitors (MkPs). Instead, fetal Mks (Kit^neg^ CD41^+^) differentiate from an immature Kit^+^ Mk intermediate.

### Yolk sac progenitors are the main source of megakaryocytes in the fetal liver until E14.5

Since fetal blood and immune cells are produced from sequential and overlapping progenitor waves (**Fig. 1E**), we next investigated the contribution of primitive cells, EMPs, and HSPCs to fetal Mks by lineage tracing using complementary Cre drivers.

The contribution of primitive cells was assessed indirectly, as no specific Cre driver targets them. We took advantage of the constitutively active *Csf1r^iCre^* strain, in which all definitive progenitors (EMP and HSPC) and mature myeloid cells were targeted. The contribution of yolk sac definitive progenitors (EMP) to Kit^+^ Mks was first detected at E9.0, while Kit^neg^ Mks were labeled later at E10.5 (**Fig. 1F**). This indicates that the primitive wave was the main source of Kit^neg^ Mks until E10. Notably, this also supported the developmental trajectory from Kit^+^ Mks to Kit^neg^ Mks.

EMP contribution to mature Mks was confirmed in pulse-labeling experiments using inducible *Cdh5^CreERT2^* animals. EMPs pulsed at E7.5 with 4-hydroxytamoxifen (4-OHT) were the main source for Kit^+^ and Kit^neg^ Mks until E14.5 and E15.5, respectively. Thereafter, HSPCs (pulsed at E10.5 with 4-OHT in *Cdh5^CreERT2^* pregnant dams) took over Mk production.

In summary, lineage tracing mouse models demonstrated that (i) the primitive wave is the main source of Mks until E10 in the yolk sac, (ii) definitive EMPs are the primary origin of Mks until late gestation (E14.5/15.5), and (iii) HSPCs supersede Mk production thereafter.

### Transcriptional regulation of megakaryopoiesis differs between stages

Next, we wanted to understand whether the stage- and ontogeny-specific Mk commitment trajectories are transcriptionally differentially regulated. To test this, we performed droplet-based paired single-cell RNA and ATAC sequencing (Chromium). We isolated cells separately at E10.5, E12.5, and E16.5 (**Fig. 2A**).

**Figure 2.**
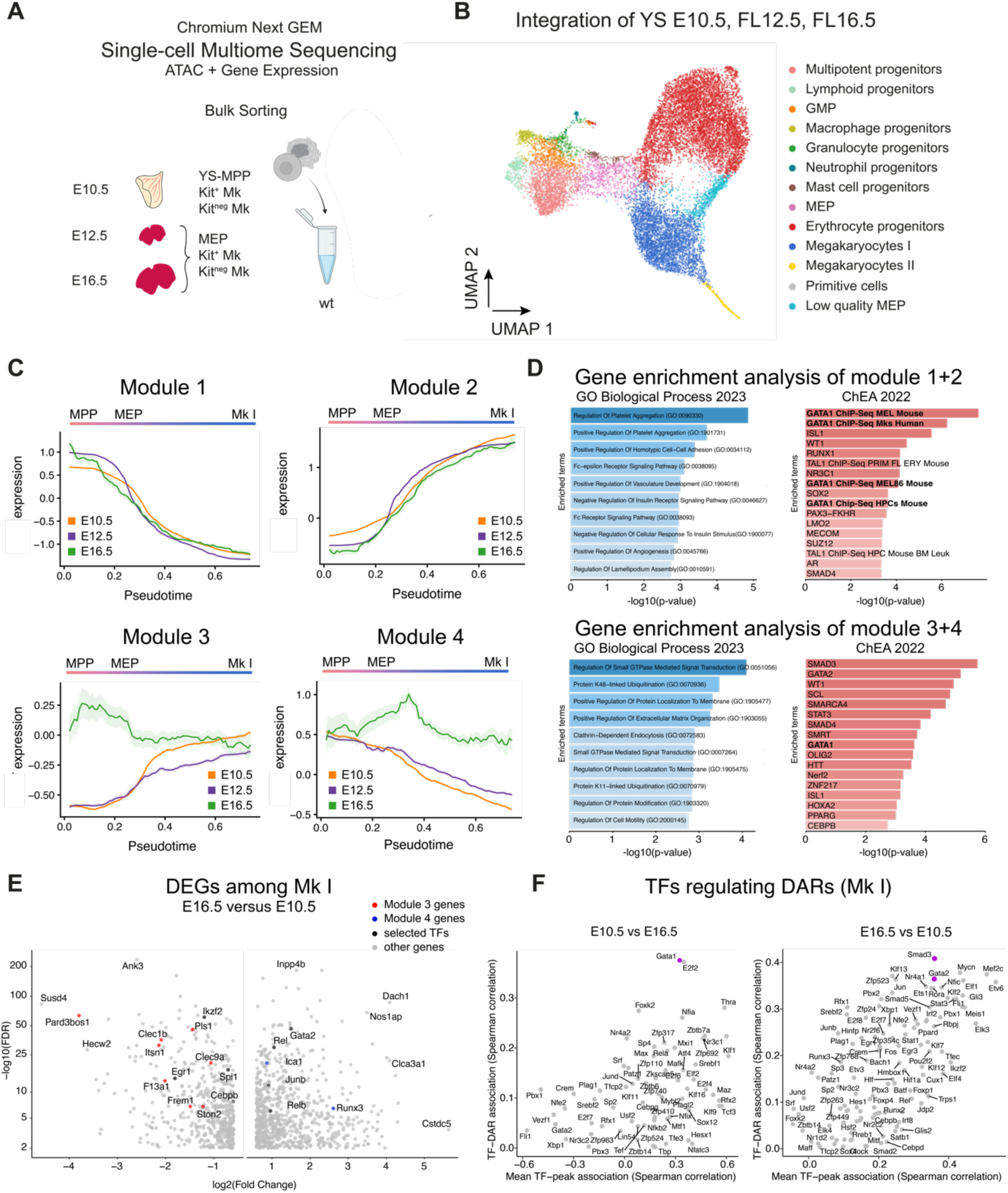
Single-cell multiome sequencing captures cells throughout megakaryopoiesis in fetal development. (**A**) Scheme illustrating sorting and sequencing strategy. Multipotent progenitors (Ter119^neg^ F4/80^neg^ Kit^high^ CD41^int^), Kit^+^, and Kit^neg^ Mks were sorted from E10.5 yolk sacs. Megakaryocyte-erythrocyte progenitors (MEP), Kit^+^, and Kit^neg^ Mks were sorted from E12.5 and E16.5 fetal livers. Wild-type (male and female) cells were sorted separately at each time point. Cells were lysed and gene expression and ATAC libraries were prepared from nuclei following the Chromium Next Gem Single-cell Multiome Sequencing pipeline. (**B**) UMAP illustrating the integration of the single-cell RNA sequencing data from E10.5 yolk sacs, E12.5, and E16.5 fetal livers. Each color labels a different cluster. GMP = granulocyte-monocyte progenitor, MEP = megakaryocyte-erythrocyte progenitor. (**C**) Graphs showing expression of co-expression modules 1 and 2 (top) and 3 and 4 (bottom) along the pseudotime from multipotent progenitors (MPP) via megakaryocyte-erythrocyte progenitors (MEP) to megakaryocytes I (Mk I) in E10.5 wild-type yolk sacs (orange), E12.5 (purple) and E16.5 (green) wild-type fetal livers. (**D**) Gene enrichment analysis (GO Biological Process 2023 (left), ChEA 2022 (right)) of genes from module 1+2 (top) or module 3+4 (bottom). (**E**) Volcano plot showing the up- and down-regulated genes in E16.5 Mk I compared to E10.5 Mk I. Module 3 and 4 genes are highlighted in red or blue, respectively. Select transcription factors are highlighted in black. (**F**) Plot representing the transcription factors (directly or indirectly) responsible for the opening of differentially accessible regions at E10.5 (left) or E16.5 (right) compared to E16.5 or E10.5, respectively, among Mk I. <u>x-axis</u>: mean of the TF-peak associations (of all differentially accessible regions at E10.5 or E16.5, respectively; Spearman correlation) indicating the likelihood that a specific TF is (directly or indirectly) responsible for the opening of a differentially accessible region. (Mean of all dots seen in supplementary data figure 4B). <u>y-axis</u>: TF-DAR association (Spearman correlation) representing the correlation of the false discovery rate of the differential accessibility of the peak at E10.5 compared to E16.5 (left) or vice versa (right) and the likelihood (TF-peak association) that a specific TF will be responsible for the opening of a differentially accessible peak. (correlation factor rho of y- and x-axis in Supplementary Data Fig. 4B). See also Supplemental Data Fig. 2, 3, 4 and Supplemental Tables 1-4.

We sequenced nuclei from 10,460 wild-type Ter119^neg^ F4/80^neg^ Kit^high^ CD41^int^ and Kit^+/neg^ CD41^+^ cells from a pool of 15 E10.5 yolk sacs (**Supplemental Data Fig. 2A**). From E12.5 (7,687 nuclei from 3 wild-type fetal livers) (**Supplemental Data Fig. 2B**) and E16.5 fetal livers (9,786 nuclei from 5 wild-type fetal livers), we sequenced nuclei from Lin^neg^ (Ter119, F4/80, Gr1, Nk1.1, CD3, CD19) Sca-1^neg^ Kit^high^ CD45^+^ CD16/32^neg^ CD34^neg^ (MEP) and Kit^+/neg^ CD41^+^ Mks (**Supplemental Data Fig. 2C**).

Cells from all three stages were integrated, clustered, and visualized using Uniform Manifold Approximation and Projection (UMAP) for dimensional reduction (**Fig. 2B, Supplemental Data Fig. 3A**). We identified multipotent progenitors, lymphoid progenitors, granulocyte-monocyte progenitors (GMPs), macrophage progenitors, granulocyte progenitors, neutrophil progenitors, mast cell progenitors, megakaryocyte-erythrocyte progenitors (MEPs), erythrocyte progenitors, 2 clusters of Mks (Mk I, Mk II), and primitive megakaryocyte-erythroblast cells (**Fig. 2B, Supplemental Data Fig. 3B**). Average expression of key marker genes for each cluster are shown in **Supplemental Data Fig. 3C**. Multipotent progenitors comprised erythro-myeloid progenitors at E10.5, hematopoietic stem and progenitor cells at E16.5 and a mixture of both at E12.5 (**Supplemental Data Fig. 3D**).

**Figure 3.**
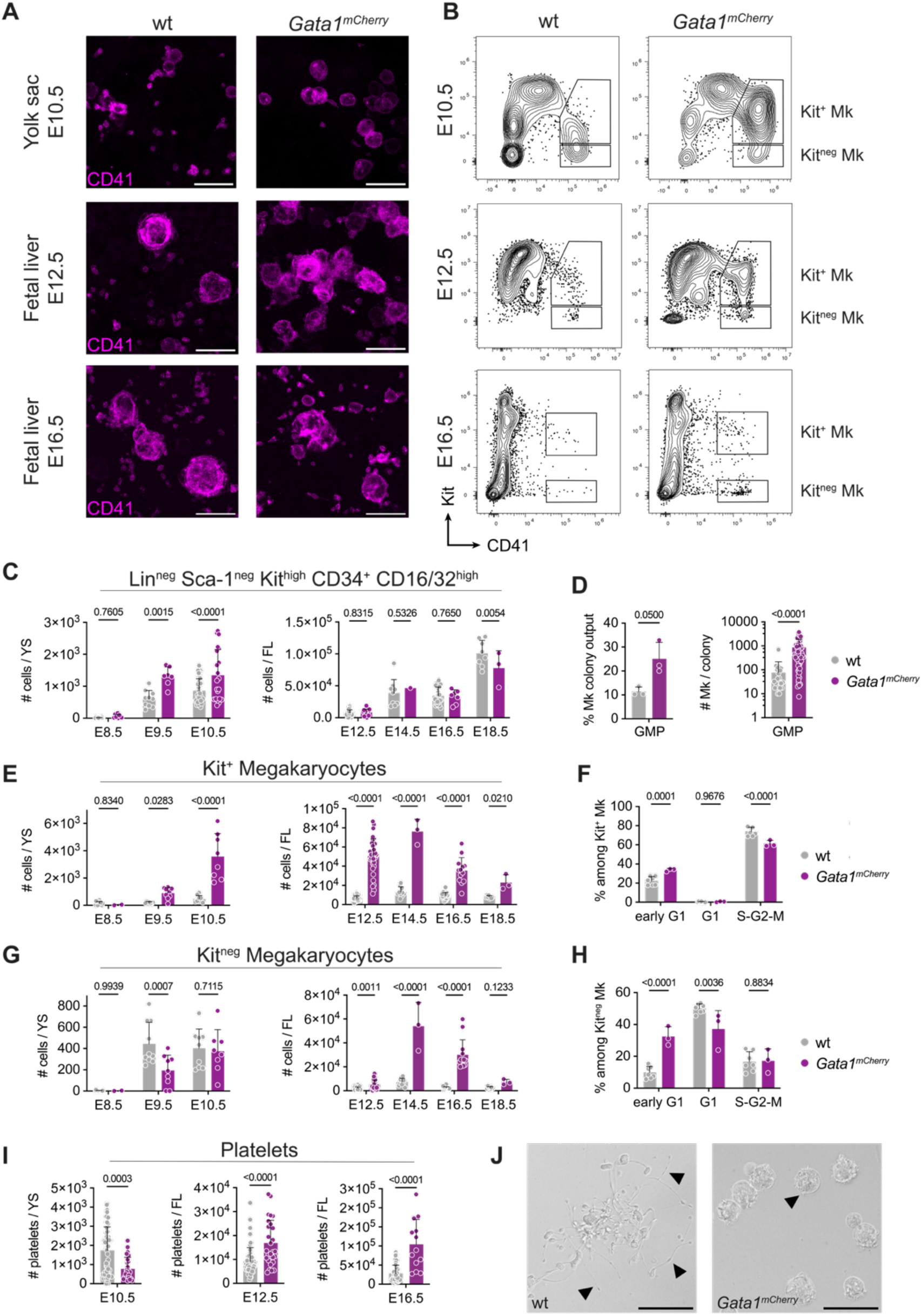
*Gata1^mCherry^* causes stage-specific accumulation of Megakaryocytes. (**A**) Immunofluorescence of 10 µm-thick maximum projection of E10.5 yolk sacs (top), 21 µm-thick maximum projection of E12.5 (middle), and 42 µm-thick maximum projection of E16.5 (bottom) fetal livers from wild-type (wt, left) and *Gata1^mCherry^* (right) embryos. Mks are stained with anti-CD41 PE (magenta). Scale bar represents 25 µm. (**B**) Dot plots showing Kit^+^ and Kit^neg^ Mk gates among Lin^neg^ cells from E10.5 yolk sac (top), Kit^+^ and Kit^neg^ Mk gates among Lin^neg^ Sca-1^neg^ cells from E12.5 (middle), and E16.5 (bottom) fetal liver from wt (left) and *Gata1^mCherry^* (right) embryos. (**C, E, G**) Bar plots showing the number of (**C**) Lin^neg^ Kit^+^ CD34^+^ CD16/32^high^ cells (immunophenotypic EMPs in the yolk sac and GMPs in the fetal liver), (**E**) Kit^+^ Mks and (**G**) Kit^neg^ Mks in E8.5-E10.5 yolk sacs (left) and E12.5-E18.5 fetal livers (right) from wt (grey) and *Gata1^mCherry^* (purple) embryos. Two-way ANOVA with Tukey’s multiple comparisons: Embryonic stage <0.0001 (C left) <0.0001 (C right) <0.0001 (E left) <0.0001 (E right) <0.0001 (G left) <0.0001 (G right); Genotype <0.0001 (C left) = 0.3267 (C right) <0.0001 (E left) <0.0001 (E right) = 0.0713 (G left) <0.0001 (G right) Interaction = 0.0423(C left) = 0.0514 (C right) <0.0001 (E left) (E right) = 0.0517 (G left) <0.0001 (G right); (**D**) Bar plot showing the frequency of colonies containing Mk progeny (Mk colony output, left) and size of Mk-containing colonies (right) from sorted single immunophenotypic GMPs grown in liquid cultures for 7 days from E12.5 fetal livers from wild-type (grey) or *Gata1^mCherry^* (purple) embryos. 3 independent litters. One-tailed Mann-Whitney test. (**F, H**) Bar plots showing the frequency of wt (grey) and *Gata1^mCherry^* (purple) Kit^+^ (**F**) and Kit^neg^ (**H**) Mks in early G1 (Fucci double negative), G1 (mCherry-Cdt^+^) and S-G2-M (mVenus-hGem^+^) phase in E12.5 fetal livers from *PGK^Cre^ R26^Fucci2aR^* embryos. n ≥3 embryos from N ≥3 litters per condition. Two-way ANOVA with Tukey’s multiple comparisons: Cell cycle: P-value <0.0001 (F) <0.0001 (H); Genotype: P-value = 0.7736 (F) = 0.1552 (H); Interaction: P-value <0.0001 (F) <0.0001 (H). (**I**) Bar plots showing the numbers of platelets in the E10.5 yolk sacs (left), E12.5 (middle), and E16.5 (right) fetal livers in wt (grey) and *Gata1^mCherry^*(purple) embryos. n >3 embryos from N >3 litters per condition. One-tailed Mann-Whitney test. (**J**) Bright-field images of Mks grown from single immunophenotypic GMPs from E12.5 fetal livers from wild-type (grey) or *Gata1^mCherry^* (purple) embryos. Scale bar represents 50 µm. Data are represented as mean ± SD. See also Supplemental Data Fig. 5.

Cells were ordered in pseudotime and we performed co-expression analysis along the Mk-differentiation trajectory from multipotent progenitors (MPPs) over MEPs to Mks (**Fig. 2C, Supplemental Data Fig. 4**). There was a core program of down-(module 1) and up-(module 2) regulated genes which was expressed at all three time points. Progenitor-specific genes such as *Gata2*, *Rac2*, *Nrip1*, *Cdk19*, *Cacnb2*, *Pstpip2, Emilin2,* and *Ptpre* were downregulated with increasing commitment and differentiation (module 1). Simultaneously, megakaryocyte-specific genes, such as *Mpl, Gucy1a1, Bin2*, *Srgap3*, and *Mmrn1,* were upregulated (module 2). Gene enrichment analysis using Enrichr (Chen et al., 2013; Kuleshov et al., 2016; Xie et al., 2021) predicted module 1 and 2 genes to be involved in platelet functions and to be predominantly controlled by GATA1 (**Figure 2D, Supplemental Table 1, 2**).

**Figure 4.**
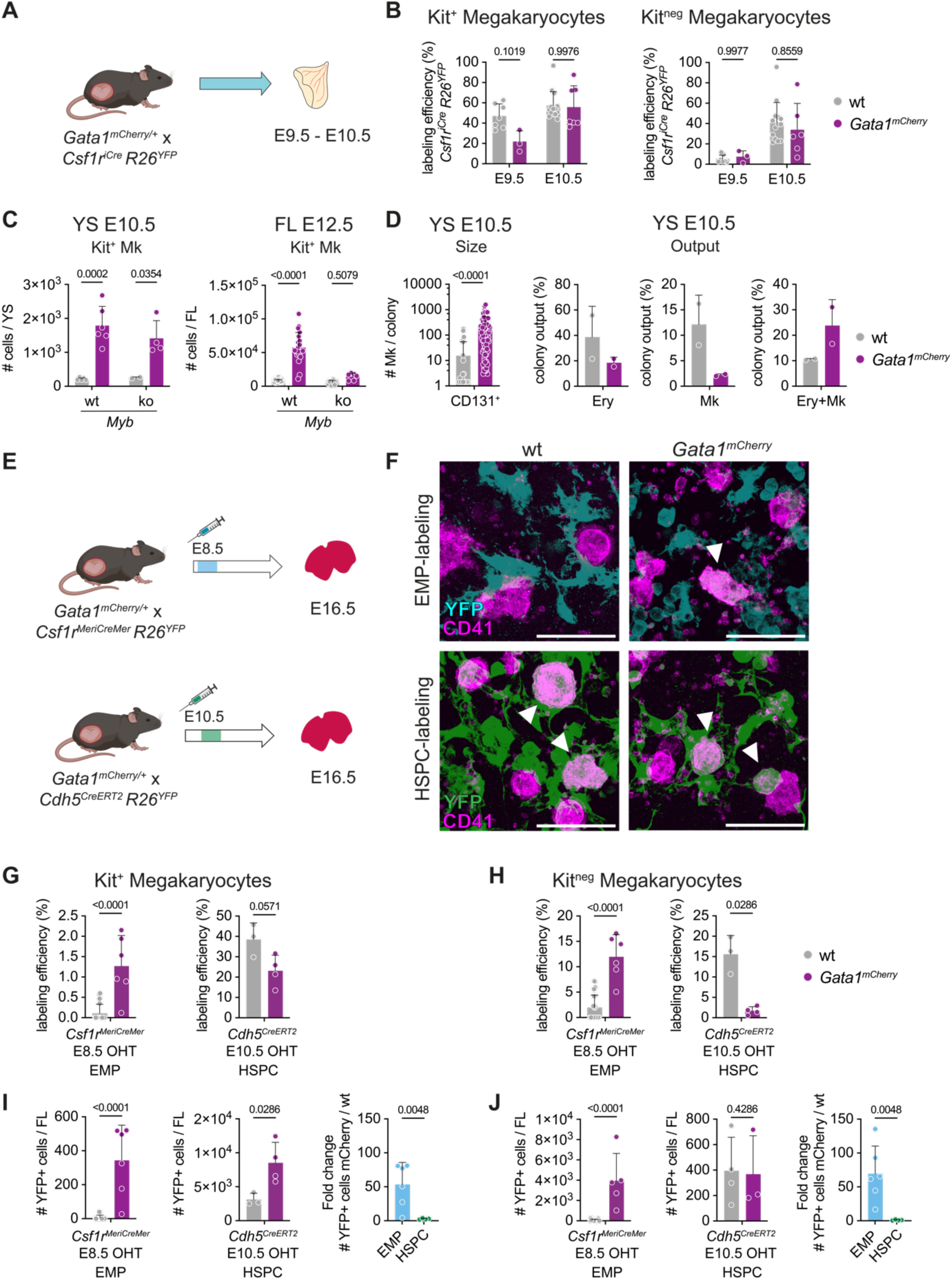
Primitive and EMP-but not HSPC-derived megakaryocytes accumulate in the presence of *Gata1^mCherry^*. (**A**) Scheme illustrating fate mapping of yolk sac-derived definitive *Gata1^mCherry^* cells in the yolk sac. (**B**) Bar plot showing labeling efficiency of Kit^+^ (left) and Kit^neg^ (right) Mks in E9.5 and E10.5 yolk sacs from wt (grey) or *Gata1^mCherry^* (purple) embryos. n ≥ 3 embryos from N ≥ 3 litters per condition. Two-way ANOVA with Tukey’s multiple comparisons: Embryonic Stage: P-value = 0.0016 (left), = 0.0004 (right); Genotype: P-value = 0.0457 (left), = 0.7610 (right); Interaction: P-value = 0.0702 (left), = 0.5466 (right). (**C**) Bar plot showing the number of Kit^+^ Mks in E10.5 yolk sacs (left) or E12.5 fetal livers (right) from *Myb^+/+^* or *Myb^-/-^ Gata1^wild-type^* (grey) or *Gata1^mCherry^* (purple) embryos. n ≥ 3 embryos from N ≥ 3 litters per condition. One-way ANOVA with Tukey’s multiple comparisons: Myb: P-value = 0.4646 (left), <0.0001 (right); Gata1: P-value <0.0001 (left), <0.0001 (right); Interaction: P-value = 0.3842 (left), <0.0001 (right). (**D**) Bar plot showing the colony size of Mk-containing colonies (far left, One-tailed Mann-Whitney test) and colony output of pure erythrocytes (Ery, left), Mks (right), and Ery+Mk mixed (far right) colonies grown from Kit^high^ CD41^+^ CD131^+^ cells from E10.5 yolk sacs. 2 independent experiments. (**E**) Scheme illustrating fate mapping of EMP-derived (top) and HSPC-derived (bottom) cells in the E16.5 fetal liver. (**F**) Immunofluorescence of 42 µm-thick maximum projection of E16.5 fetal livers from wild-type (wt, left) and *Gata1^mCherry^* (right) embryos. Mks are stained with anti-CD41 PE (magenta). EMP-derived cells are fate-mapped in *Csf1r^MeriCreMer^ R26^YFP^* embryos pulsed at E8.5 with 4-OHT (cyan). HSPC-derived cells are fate-mapped in *Cdh5^CreERT2^ R26^YFP^* embryos pulsed at E10.5 with 4-OHT (green). Scale bar represents 50 µm. White arrow heads indicate YFP^+^ Mks. (**G, H**) Bar plot showing labeling efficiency of EMP-derived (left) and HSPC-derived (right) Kit^+^ (**G**) and Kit^neg^ (**H**) Mks in E16.5 fetal livers from wild-type (grey) or *Gata1^mCherry^* (purple) embryos. n ≥ 3 embryos from N ≥ 3 litters per condition. One-tailed Mann-Whitney test. (**I, J**) Bar plots showing the number of EMP-(left) and HSPC-derived (middle) YFP^+^ Kit^+^ (**I**) and Kit^neg^ (**J**) Mks in E16.5 fetal livers from wild-type (grey) or *Gata1^mCherry^*(purple) embryos. Fold change (right) of *Gata1^mCherry^* over wild-type EMP-(blue) and HSPC-derived (green) Kit^+^ (**I**) and Kit^neg^ (**J**) Mks. n ≥ 3 embryos from N ≥ 3 litters per condition. One-tailed Mann-Whitney test. Data are represented as mean ± SD. See also Supplemental Data Fig. 6.

There were also significant differences in transcription between megakaryopoiesis at E10.5/E12.5 and E16.5. The expression of module 3 genes (*Frem1*, *F13a1, Plxna4, Clec1b, Arhgap18, Arhgap45,* etc.) was stable throughout megakaryopoiesis at E16.5, while at E10.5 and E12.5, they were upregulated after the MEP stage. In a mirror image, the expression of module 4 genes, such as *Ica1*, *Zmiz1*, *Runx3*, and *Adgrg3*, was sustained throughout Mk differentiation at E16.5, but was downregulated at E10.5 and E12.5 (**Fig. 2C, Supplemental Data Fig. 4A**). Following the stage-dependent dynamics of the expression of module 3 and 4 genes, E16.5 Mks expressed higher levels of module 4 genes and lower levels of module 3 genes compared to E10.5 Mks (**Fig. 2E**). Module 3 and 4 genes are involved in the regulation of GTPase-mediated signaling and play a role in the regulation of the cell membrane (**Figure 2D, Supplemental Table 3, 4**). A set of these genes (e.g. *Smad3*, *Zmiz1*, *Rasgef1b*, *Tmem40*, *Itsn1*, *Dock8*) were predicted to be regulated by a group of transcription factors, including SMAD3, SMAD4, WT1, SCL, LMO2, GATA2, and also GATA1. Furthermore, HOXA2, which modulates only yolk sac-derived and not adult Mks (Iacovino et al., 2007), was predicted to regulate genes (*Arhgap18*, *Frem1*) that were expressed at higher levels at E10.5 than at E16.5 (**Fig. 2D, E**).

Among the transcription factors predicted to regulate module 3 and 4 genes, GATA1 was one of the most likely to regulate the differentially accessible regions at E10.5. By contrast, SMAD3 and GATA2 were among the transcription factors responsible for the differentially accessible regions at E16.5 (**Fig. 2F, Supplemental Data Fig. 4B**). This suggests that GATA1 may not only have common functions in megakaryopoiesis across stages but also ontogeny-specific roles.

Altogether, yolk sac and intraembryonic progenitors follow distinct trajectories at the transcriptional and cellular level to make Mks. In this process, GATA1, a key transcription factor in megakaryopoiesis regulating Mk commitment, terminal maturation, and platelet formation (Shivdasani et al., 1997; Iwasaki et al., 2003), has both common (module 1+2) but also ontogeny-specific (module 3+4) functions in yolk sac-derived versus HSPC-derived Mk development.

### *Gata1^mCherry^* causes stage-specific accumulation of Megakaryocytes

These ontogeny-specific features of GATA1 during fetal hematopoiesis might be a driving factor of the stage-specific effects of *Gata1* mutations (Shivdasani et al., 1997; Shimizu et al., 2001; Li et al., 2005). In our hands, expression of *Gata1^mCherry^,* encoding for a fusion protein of GATA1 and the fluorescent protein mCherry (Hoppe et al., 2016), caused a stage-specific accumulation of Mks in the yolk sac and the fetal liver in both male and female murine embryos (**Fig. 3A, B, Supplemental Data Fig. 5a**). No accumulation of megakaryocyte progenitors or Mks was detected in the bone marrow of 6-month-old mice (**Supplemental Data Fig. 5B, C**).

**Figure 5.**
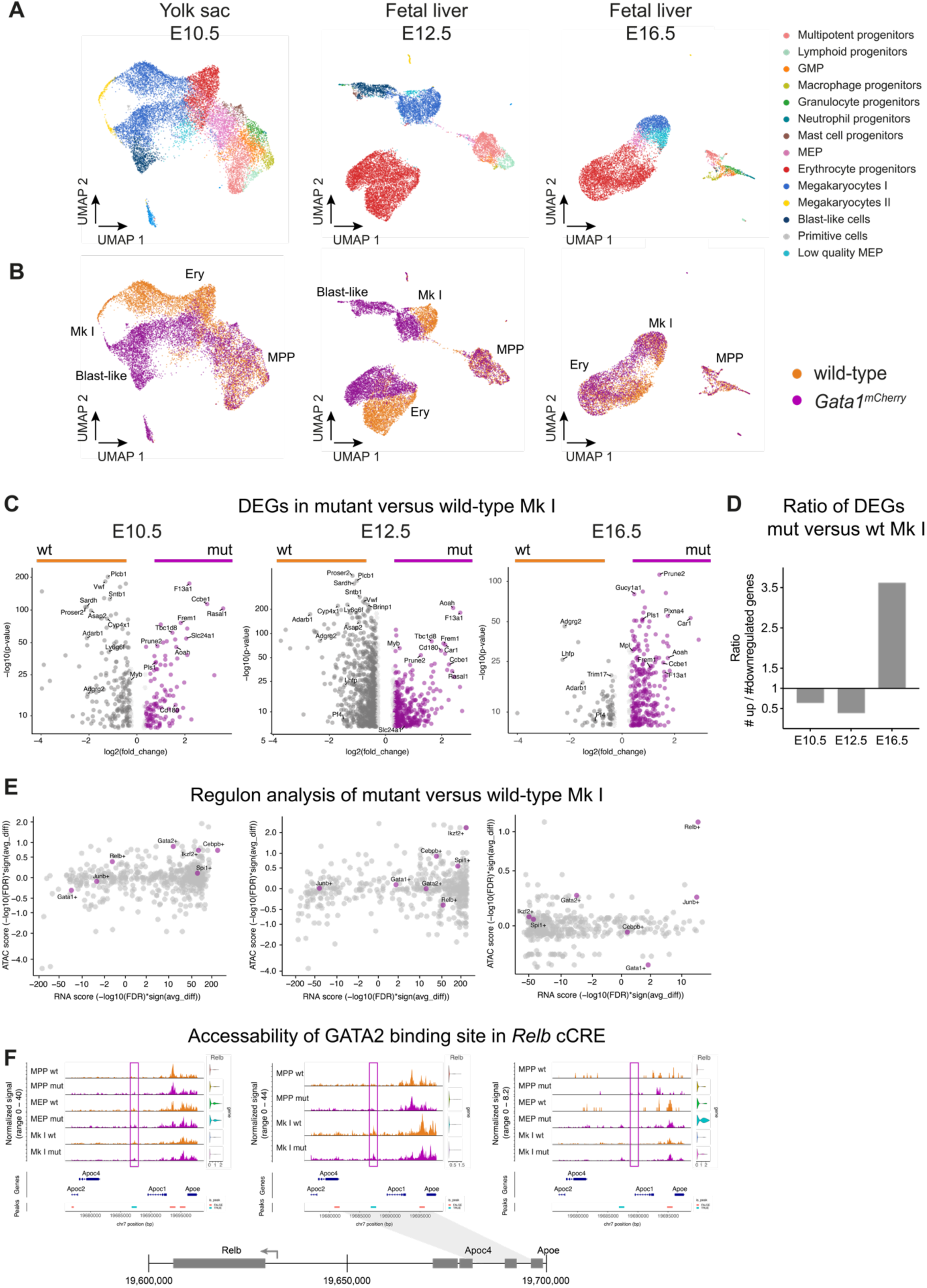
*Gata1^mCherry^* drives stage-specific transcriptional changes in megakaryopoiesis. (**A**) UMAPs from E10.5 yolk sacs, E12.5, and E16.5 fetal livers featuring clusters identified in the integration of the single-cell RNA sequencing data of all three time points together. (**B**) UMAPs featuring wild-type (orange) and *Gata1^mCherry^* (purple) cells in single-cell RNA sequencing datasets from E10.5 yolk sacs, E12.5, and E16.5 fetal livers. (**C**) Volcano plot showing up (purple) and down (grey) regulated genes in *Gata1^mCherry^*Mks (MkI, purple bar) compared to wild-type (grey bar). (**D**) Bar plot quantifying ratio of up versus down regulated genes between wild-type and *Gata1^mCherry^* Mks (Mk I) at E10.5, E12.5 and E16.5. (**E**) RNA- and ATAC scores calculated by Pando analysis indicating the increase in transcription factor activity in *Gata1^mCherry^* compared to wt Mks (Mk I) in E10.5 yolk sacs, E12.5, and E16.5 fetal livers. (**F**) Coverage plot illustrating chromatin accessibility of GATA2 binding site in a *Relb* candidate cis-regulatory element (cCRE). See also Supplemental Data Fig. 7.

To identify the stages of megakaryopoiesis in which *Gata1^mCherry^*exerts its effect, we characterized Lin^neg^ Kit^high^ Sca-1^neg^ CD34^+^ CD16/32^high^ progenitors (named EMP in the yolk sac, GMP in the fetal liver (Akashi et al., 2000; McGrath et al., 2015)), Mks (Kit^+^ and Kit^neg^) and platelets in *Gata1^mCherry^* mutant embryos. We focused on the two main fetal hematopoietic niches, the yolk sac (from E8.5 to E10.5) and fetal liver (from E12.5 to E18.5).

First, while Lin^neg^ Kit^high^ Sca-1^neg^ CD34^+^ CD16/32^high^ progenitors were modestly increased in the yolk sac, their numbers were unaffected in the fetal liver (**Fig. 3C**). However, when cultured, fetal liver Lin^neg^ Kit^high^ Sca-1^neg^ CD34^+^ CD16/32^high^ progenitors had a higher Mk output and yielded larger-sized Mk colonies (**Fig. 3D, Supplemental Data Fig. 5D**). Thus, *Gata1^mCherry^* caused hyperproliferation and Mk-bias downstream of Lin^neg^ Kit^high^ Sca-1^neg^ CD34^+^ CD16/32^high^ progenitors.

Accordingly, Kit^+^ Mks were affected by *Gata1^mCherry^*. Accumulation of Kit^+^ Mks started in the yolk sac at E9.5 (5-fold increase), peaked around E12.5 in the fetal liver (7-fold increase), and decreased thereafter (**Fig. 3E, Supplemental Data Fig. 5B**). In a Fucci mouse model, in which cells in G1 and S-G2-M can be distinguished by the presence of mCherry-Cdt or mVenus-hGem fusion proteins, respectively, no overt differences in the cell cycle of Kit^+^ Mks were found between wild-type and *Gata1^mCherry^*embryos (**Fig. 2F**) (note that mCherry-Cdt is brighter than *Gata1^mCherry^* which allows to distinguish them easily by flow cytometry). However, only *Gata1^mCherry^* but not wild-type Kit^+^ Mks showed colony forming potential (**Supplemental Data Fig. 5E**).

Numbers of Kit^neg^ Mks, by contrast, were reduced at E9.5, unchanged at E10.5 in the yolk sac, and increased 6-fold at E14.5 in the fetal liver (**Fig. 3G, Supplemental Data Fig. 5C**). This accumulation was transient and resolved afterwards (**Supplemental Data Fig. 5C**). The frequency of *Gata1^mCherry^* Kit^neg^ Mk in S-G2-M was unaffected (**Fig. 3H**), suggesting this accumulation was not due to self-amplification of the Kit^neg^ population but rather due to increased input from their upstream precursors, the Kit^+^ Mks, and decreased output by blocked maturation into platelet-producing cells. Indeed, platelet numbers were reduced in the yolk sac at E10.5 but were increased in the fetal liver at E12.5 and E16.5 (**Fig. 3A, I**). This delay in platelet production was in line with the finding that *Gata1^mCherry^* Mks did not form proplatelets like wild-type Mks after 7 days of incubation in liquid cultures (**Fig. 3J**).

Altogether, our findings show that *Gata1^mCherry^* has stage-specific effects. Between E9.5 and E14.5, *Gata1^mCherry^*blocks maturation downstream of Lin^neg^ Kit^high^ Sca-1^neg^ CD34^+^ CD16/32^high^ progenitors, causing an abnormal accumulation of Mks and a delayed production of platelets.

### Primitive and EMP-but not HSPC-derived megakaryocytes accumulate in the presence of *Gata1^mCherry^*

The stage-specific accumulation of Mks driven by *Gata1^mCherry^*coincided with the ontogeny switch from EMP to HSPC seen in wild-type embryos. Therefore, we tested whether the stage-specific phenotype of *Gata1^mCherry^* was ontogeny-dependent.

First, we crossed the *Gata1^mCherry^* allele with the *Csf1r^iCre^* fate-mapping model described before (**Fig. 4A**) and found that primitive Kit^+^ Mks accumulated at E9.5 in *Gata1^mCherry^* yolk sacs. In contrast, Kit^+^ and Kit^neg^ Mks accumulating at E10.5 were EMP-derived, as in control embryos (**Fig. 4, Supplemental Data Fig. 6A**).

**Figure 6.**
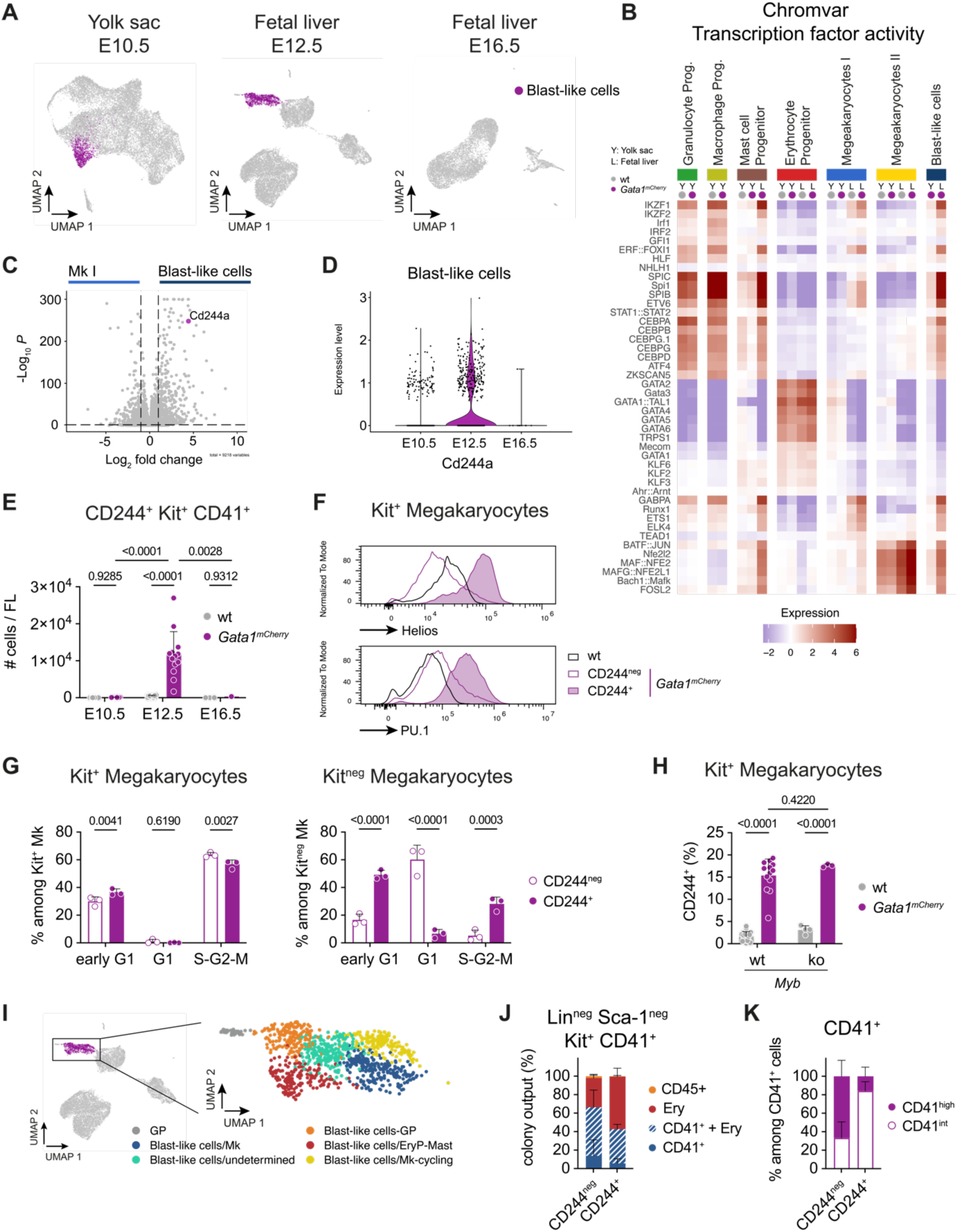
*Gata1^mCherry^*-driven formation of CD244^+^ blast-like cells occurs exclusively in yolk sac-derived megakaryopoiesis. (**A**) UMAPs featuring blast-like cells (purple) in the E10.5 yolk sac, E12.5, and E16.5 fetal liver. (**B**) Heat map showing transcription factor activity calculated using Chromvar among clusters in yolk sac (Y) and fetal liver (L) wild-type (grey) and *Gata1^mCherry^* (purple) samples. (**C**) Volcano plot comparing the gene expression between mutant blast-like cells and wild-type and mutant Mks (Mk I) in the E12.5 fetal liver. *Cd244a* is highlighted in purple. (**D**) Violin plot showing the expression level of *CD244a* among blast-like cells at E10.5, E12.5, and E16.5. (**E**) Bar plot showing the number of CD244^+^ Kit^+^ CD41^+^ cells per E12.5 fetal liver from wild-type (grey) or *Gata1^mCherry^* (purple) embryos. ≥ 3 embryos from ≥ 3 litters per condition except for E16.5 *Gata1^mCherry^*(only 1 embryo). Two-way ANOVA: Embryonic Stage: P-value <0.0001. Genotype: P-value = 0.0052. Interaction: P-value <0.0001. (**F**) Histograms showing Helios (top) and PU.1 (bottom) protein expression in wild-type (black), *Gata1^mCherry^* CD244^neg^ (purple border), and *Gata1^mCherry^* CD244^+^ (purple filled) cells from E12.5 fetal livers. (**G**) Bar plot showing the frequency of CD244^neg^ (purple border) and CD244^+^ (purple filled) Kit^+^ (left) and Kit^neg^ (right) Mks in early G1 (Fucci double negative), G1 (Fucci mCherry^+^) and S-G2-M (Fucci mVenus^+^) phase in E12.5 fetal livers from embryos with constitutive expression of *R26^Fucci/+^*. n ≥3 embryos from N ≥3 litters. Two-way ANOVA with Tukey’s multiple comparisons. Cell cycle: P-value <0.0001 (left), = 0.0003 (right); Population: P-value = 0.6747 (left), = 0.8054 (right); Interaction: P-value = 0.0009 (left), P-value <0.0001 (right); (**H**) Bar plot showing the frequency of CD244 expression among Kit^+^ Mks from *Myb^+/+^*or *Myb^-/-^ Gata1^wild-type^* (grey) or *Gata1^mCherry^*(purple) embryos. ≥ 3 embryos from ≥ 3 litters per condition. Two-way ANOVA. Gata1: P-value <0.0001; Myb P-value = 0.0585; Interaction: P-value = 0.6915. (**I**) UMAP of blast-like cells from E12.5 fetal livers featuring different subpopulations: GP = granulocyte progenitors, EryP = erythrocyte progenitor, Mk = megakaryocyte. (**J**) Stacked bar plot illustrating the colony output of CD244^neg^ and CD244^+^ Kit^+^ Mks from *Gata1^mCherry^* E12.5 fetal livers. Detected were CD45^+^ cells, erythrocytes (Ery, CD71^+^, Ter119^+^ cells), and CD41^+^ cells. 3 independent experiments. (**K**) Stacked bar plot showing the frequency of CD41^high^ and CD41^int^ cells among all CD41^+^ (pure CD41^+^ and CD41^+^ + Ery) cells detected in **J**. Data are represented as mean ± SD. See also Supplemental Data Fig. 8, 9.

Recently, we discovered that EMPs give rise to two distinct waves of Mks that can be distinguished by their spatiotemporal kinetics and dependency on the transcription factor c-*Myb* (Tober, McGrath & Palis, 2008; Iturri et al., 2021). Lack of *Myb* does not hamper direct Mk differentiation from EMPs in the yolk sac. However, in *Myb*-deficient embryos EMPs fail to produce bipotent CD131^+^ erythro-Mk progenitors causing a lack of erythroid and Mk generation in the fetal liver (**Supplemental Data Fig. 6B**).

We thus examined Mk numbers in double *Gata1^mCherry^ Myb*-deficient yolk sacs and fetal livers. Mks accumulated normally in double *Gata1^mCherry^ Myb* mutant yolk sacs, confirming that *Gata1^mCherry^* affects the direct differentiation of Mks from EMPs (**Fig. 4C**). In contrast, Mk accumulation was abrogated in double *Gata1^mCherry^ Myb^-/-^* fetal livers (**Fig. 4C**). This demonstrated that Mks accumulating in *Gata1^mcherry^* fetal livers arise from EMP-derived *Myb*-dependent CD131^+^ progenitors.

To test the effect of *Gata1^mcherry^* in the production of Mks from CD131^+^ progenitors, we performed colony-forming assays. *Gata1^mCherry^* CD131^+^ progenitors gave rise to more mixed colonies with erythrocyte and Mk output at the expense of pure erythroid and pure Mk colonies. Further, colonies containing Mks were larger in *Gata1^mCherry^* conditions than in wild-type conditions (**Fig. 4D, Supplemental Data Fig. 6C**). Altogether, primitive and both EMP-derived Mk pathways are affected by *Gata1^mCherry^* and contribute to Mk accumulation in the yolk sac and fetal liver.

To specifically trace the fate of *Gata1^mCherry^* EMPs and HSPCs in the fetal liver, we pulse-labeled EMPs at E8.5 in inducible *Gata1^mCherry^ Csf1r^MeriCreMer^ R26^eYFP^* embryos and HSPCs at E10.5 in *Gata1^mCherry^ Cdh5^CreERT2^ R26^eYFP^* embryos (**Fig. 4E**). EMP-derived Mks continued to accumulate until E16.5 (**Fig. 4F**). Increased labeling efficiency in both Mk subsets in *Gata1^mCherry^* EMP fate mapped (E8.5 OHT in *Csf1r^MeriCreMer^ R26^eYFP^*) embryos associated with the decreased labeling efficiency in *Gata1^mCherry^*HSPC fate mapped (E10.5 OHT in *Cdh5^CreERT2^ R26^eYFP^*) embryos demonstrated that accumulating Mks in *Gata1^mCherry^* fetal livers were moslty of EMP origin (**Fig. 4G, H**).

In contrast to wild-type conditions where the majority of Mks are HSPC-derived at E16.5, Kit^neg^ Mks in *Gata1^mCherry^* embryos were still of EMP origin. Importantly, when analyzing the number of pulse-labeled cells (**Fig. 4I, J**), HSPC-traced Kit^+^ Mks did accumulate in the fetal liver at E16.5, albeit at a lower extent than EMP-derived ones. Indeed, while the number of EMP-derived Kit^+^ and Kit^neg^ Mks showed a 50-fold increase, HSPC-derived Kit^+^ Mks only increased in numbers by 2-fold, and numbers of HSPC-derived Kit^neg^ Mks were unchanged (**Fig. 4I, J**).

Collectively, these data suggest that only yolk sac-derived (primitive and EMP) and not intraembryonic HSPC-derived megakaryocytes strongly accumulate in the presence of *Gata1^mCherry^*. Moreover, analysis of double *Gata1^mCherry^ Myb* mutants provides evidence that the phenotype exerted by *Gata1^mCherry^* is not dependent on the differentiation route but rather on the progenitor of origin.

### *Gata1^mCherry^* drives stage-specific transcriptional changes in megakaryocytes

To understand the molecular mechanisms underlying the ontogeny-specific effects of *Gata1^mCherry^,* we compared wild-type and *Gata1^mCherry^* cells using droplet-based single-cell RNA and ATAC sequencing (Chromium). We sorted *Gata1^mCherry^* (male) cells separately at E10.5, E12.5, and E16.5 (**Supplemental Data Fig. 7A**).

We sequenced nuclei 13,742 *Gata1^mCherry^* Ter119^neg^ F4/80^neg^ Kit^+^ CD41^int^ and Kit^+/neg^ CD41^+^ cells from a pool of 9 the E10.5 yolk sacs (**Supplemental Data Fig. 2A**). From E12.5 (9,208 nuclei from 4 *Gata1^mCherry^* fetal livers) (**Supplemental Data Fig. 2B**) and E16.5 fetal livers (5,858 nuclei from 4 *Gata1^mCherry^* fetal livers), we sequenced nuclei from Lin^neg^ (Ter119, F4/80, Gr1, Nk1.1, CD3, CD19) Sca-1^neg^ Kit^high^ CD45^+^ CD16/32^neg^ CD34^neg^ (MEP) and Kit^+/neg^ CD41^+^ Mks (**Supplemental Data Fig. 2C**).

Single-cell data of *Gata1^mCherry^* cells were combined with wild-type cells and clustered using Uniform Manifold Approximation and Projection (UMAP) for dimensional reduction for each stage individually. Mutant cells were found in all clusters first identified in the wild-type dataset. In addition, mutant cells also formed a cluster of blast-like cells which did not exist in wild-type samples (**Fig. 5A, B, Supplemental Data Fig. 7B**). At all three stages, wild-type (orange) and *Gata1^mCherry^*(purple) multipotent and committed myeloid and lymphoid progenitors integrated well. In contrast, wild-type and mutant cells of the Mk and erythrocyte lineage integrated at E16.5 but not at E10.5 and E12.5 (**Fig. 5B**).

A core of up-regulated genes (such as *Frem1*, *F13a1*, *Aoah*) was shared among mutant Mk I at all three time points (**Fig. 5C, Supplemental Data Fig. 7C**). In line with the stage-specific features observed in wild-type megakaryopoiesis, *Gata1^mCherry^*exerted mostly repressive functions at E10.5 and E12.5 (more down-than up-regulated genes), whereas it activated gene expression at E16.5 (predominantly up-regulated genes) (**Fig. 5C, D Supplemental Data Fig. 7C, D**).

Furthermore, regulon analysis revealed *Gata1^mCherry^*-driven changes in transcription factor activity that differed between stages (**Fig. 5E**). CEBPB was the transcription factor with the highest increase in activity among Mk I in *Gata1^mCherry^* at E10.5, Helios (*Ikzf2*) and PU.1 (*Spi1*) at E12.5 and JUNB and RELB at E16.5 (**Fig. 5E**). CEBPB showed no increase in activity at E16.5 and Helios (*Ikzf2*) and PU.1 (*Spi1*) even showed decreased activity at E16.5.

This is in line with our finding that in wild-type conditions, Mk I expressed higher levels of *Cebpb*, *Spi1*, and *Ikzf2* at E10.5, while wild-type E16.5 Mk I showed higher levels of *Gata2, Junb,* and *Relb* (**Figure 2E**). Intriguingly, a GATA2 binding site in a *Relb* candidate cis-regulatory element (cCRE) was shut down in wild-type MEPs and Mks at E16.5 but was accessible in mutant cells (**Fig. 5F**). At E10.5 and E12.5, this GATA2 binding site was similarly accessible in mutant and wild-type MEPs and Mks. GATA2 is a transcription factor that is induced in HSPCs during hematopoietic specification and interacts with Gata1 during Mk commitment and differentiation (Tsai et al., 1994; Doré & Crispino, 2011; Gao et al., 2013). It has unique regulatory features in yolk sac and fetal liver megakaryopoiesis compared to bone marrow Mk production (Fujiwara et al., 2004; Huang et al., 2009).

Taken together these data suggest that *Gata1^mCherry^* disrupts megakaryopoiesis differently and more severely at E10.5 and E12.5 compared to E16.5 at the transcriptional level.

### *Gata1^mCherry^*-driven formation of CD244^+^ blast-like cells occurs exclusively in yolk sac-derived megakaryopoiesis

In line with the stage-specific differences at the cellular level, single-cell RNA sequencing revealed a cluster of blast-like cells which were only detected exclusively in *Gata1^mCherry^* and not in wild-type embryos, and only at E10.5 and E12.5 but not at E16.5 (**Fig. 6A**).

These blast-like cells expressed typical Mk genes such as *Pf4*, *Itga2b* (CD41), *Nfe2*, and *Rap1b*, but also upregulated myeloid and pluripotency genes such as *Spi1* and *Myb* (**Supplemental Data Fig. 8A**). Likewise, blast-like cells showed similar patterns of transcription factor activity as Mks, but also abnormal activity of myeloid and pluripotency transcription factors such as *Spi1*, *Cebpa, Cebpb, Ikzf1* and *Ikzf2* at E10.5 and E12.5 (**Fig. 6B, Supplemental Data Fig. 8B**). While the activity of few transcription factors was changed at E10.5 in blast-like cells, many transcription factors had increased activity at E12.5 (**Supplemental Data Fig. 8B**).

To functionally characterize blast-like cells, we searched for cell surface markers specifically expressed in mutant blast-like cells but not wild-type or mutant Mks. *Cd244a*, a type-I transmembrane protein belonging to the signaling lymphocytic activation molecule family of receptors (SLAMF), was significantly upregulated in blast-like cells at E12.5 (**Fig. 6C**). On the contrary, *Cd244a* was barely expressed at E10.5 and absent at E16.5 (**Fig. 6D**). We validated *Cd244a* expression at the protein level using flow cytometry. CD244^+^ cells were only found in *Gata1^mCherry^* fetal livers at E12.5 *in vivo* and were present among Kit^+^ and Kit^neg^ CD41^+^ Mks (**Fig. 6E, Supplemental Data Fig. 8C, D**). CD244^+^ Kit^+^ CD41^+^ blast-like cells showed higher protein levels of Helios (*Ikzf2*) and PU.1 (*Spi1*) than CD244^-^ Kit^+^ CD41^+^ cells, confirming the upregulation of myeloid and pluripotency transcription factors from the single-cell sequencing data (**Fig. 6B, F, Supplemental Data Fig. 8A, B**).

At E10.5, blast-like cells were detected at the transcriptional but not at the cellular level. Although E10.5 blast-like cells did not express *CD244a* mRNA, the *Cd244a* promoter was accessible in blast-like cells at both E10.5 and E12.5. In contrast, *the Cd244a* promoter was closed in both wt and *Gata1^mCherry^* Mks at E10.5 and E12.5 (**Supplemental Data Fig. 8E**).

These data, together with the observation that the overall transcription factor activity in blast-like cells increases from E10.5 to E12.5 (**Supplemental Data Fig. 8B**), indicated that blast-like cell fate is primed in *Gata1^mCherry^* yolk sac progenitors. In the fetal liver, these blast-like cells then mature as evidenced by the expression of CD244, Helios, and PU.1 proteins (**Figure 6E, F**). The fact that *Gata1^mCherry^*-driven blast-like cells were transient and peaked at E12.5, suggests that blast-like cell development is restricted to yolk sac lineages and does not occur in HSPC-derived megakaryopoiesis.

### *Gata1^mCherry^*-driven blast-like cells are hyperproliferative cells that retain erythrocyte potential

Finally, we functionally characterized the fetal liver blast-like cells at E12.5. Kit^+^ CD244^+^ blast-like cells were as proliferative as Kit^+^ CD244^neg^ Mks, whereas Kit^neg^ CD244^+^ blast-like cells were more proliferative than Kit^neg^ CD244^neg^ Mks (**Fig. 6G**). Even though blast-like cells were highly proliferative, their development did not depend on *Myb* (**Fig. 6H**).

Further, sub-clustering of blast-like cells at E12.5 revealed their heterogeneity. While cells in the blast-like cell cluster expressed high levels of *Itga2b* (CD41) and *Gata2*, we identified sub-clusters with granulocyte (*Gria3*), megakaryocyte (*Mpl*), or erythrocyte-mast cell (*Tgfbr3*, *Klf1, Ifitm1*) gene expression signatures (**Supplemental Data Fig. 9A**). *CD244a* expression was limited to non-Mk clusters (**Supplemental Data Fig. 9B**).

Despite the expression of myeloid genes such as *Spi1* (PU.1) at the mRNA and protein level (**Fig. 6F & Supplemental Data Fig. 9A**), Kit^+^ CD41^+^ CD244^+^ blast-like cells did not give rise to macrophages, neutrophils, or mast cells when cultured for 7 days (**Fig. 6J**). However, CD244^+^ cells did show increased red blood cell potential compared to CD244^neg^ Kit^+^ CD41^+^ cells, which came at the expense of CD41^+^ Mk output (**Fig. 6J**). Concerning the CD41^+^ progeny of cultured Kit^+^ CD41^+^ cells, Mk potential (CD41^high^ in culture) was enriched in CD244^neg^ cells, whereas CD244^+^ cells mainly produced CD41^int^ undifferentiated cells (**Fig. 6K and Supplemental Data Fig. 9C**).

Collectively, accumulating *Gata1^mCherry^* Kit^+^ Mks had cloning capacity, in stark contrast to wild-type Kit^+^ Mks. They contained at least two subsets of cells: CD244^neg^ Mks endowed with high Mk potential and CD244^+^ blast-like cells. CD244^+^ blast-like cells were less committed towards Mk fate and instead gave rise to erythrocytes and CD41^int^ undifferentiated cells. The overexpression of myeloid transcription factors such as *Spi1* (PU.1) did not translate into myeloid differentiation *in vitro*. Rather, the expression of these transcription factors was a sign of a *Gata1^mCherry^*-driven delay in the lineage commitment towards erythrocyte and Mk fate.

## Discussion

This study reveals ontogeny-specific differences in megakaryopoiesis throughout development. Transcription factor perturbations, here expression of *Gata1^mCherry^* which serves as a model for mutant Gata1, exacerbate these ontogeny-specific differences and cause a stage-specific accumulation of immature megakaryocytes.

### Gata1^mCherry^ phenocopies Gata1s

A key finding of this study is that *Gata1^mCherry^* mice phenocopy *Gata1s (Gata1short)* mouse models (Li et al., 2005; Ling et al., 2019). Naturally, both the short and full-length isoforms of *Gata1* are expressed (Calligaris et al., 1995; Rainis et al., 2003). In *Gata1s* mice, however, somatic point mutations in exon 2 lead to the exclusive expression of the short and loss of the long isoform (Rainis et al., 2003). Both *Gata1s* and *Gata1^mCherry^* mouse models are characterized by a transient hyperproliferation and a block in maturation that ultimately lead to the accumulation of immature Mks in the yolk sac and fetal liver (Li et al., 2005; Juban et al., 2021). Additionally, erythropoiesis is delayed in *Gata1s* and *Gata1^mCherry^* embryos (data not shown for *Gata1^mCherry^)* (Ling et al., 2019).

*Gata1s* lacks the N-terminal transactivation domain. This domain does not bind DNA binding nor friend-of-GATA1 (FOG-1) (Tsang et al., 1998; Wechsler et al., 2002; Kaneko et al., 2012). The N-terminal domain is important for controlling proliferation and terminal Mk differentiation (Kuhl et al., 2005). Moreover, primitive and definitive hematopoiesis each require different dosages of the N-terminal transactivation domain (Shimizu et al., 2001).

In contrast to *Gata1s, Gata1^mCherry^* is a fusion protein of mCherry to the C-terminus of endogenous GATA1. Several non-mutually exclusive phenomena could explain why *Gata1^mCherry^* phenocopies *Gata1s*. First, mCherry might decrease Gata1 mRNA or protein stability. Importantly, *Gata1^mCherry^* animals do not display the decrease in male viability nor the neonatal anemia and thrombocytopenia observed in *Gata1^low^* mutants (Shivdasani et al., 1997; Hoppe et al., 2016), thereby supporting that the fusion of mCherry to GATA1 rather disrupts the function of GATA1 at the protein level, by compromising its 3D organization. Second, mCherry might sterically hinder the accessibility of the N-terminal domain and block the interaction with yet-to-be-identified binding partners (Elagib et al., 2003; Xu et al., 2006; Ling & Crispino, 2020). Finally, mCherry might compromise potential intramolecular interactions between the C- and T-terminal transactivating domains, thereby changing the functionality of GATA1 transcriptional activity. These functional changes could lead to increased binding strength of GATA1 to the DNA (Zambo et al., 2024), altered recruitment of C-terminal and/or N-terminal transcriptional co-activators/repressors, or aberrant chromatin accessibility (Chlon et al., 2015; Ling et al., 2019).

A key protein interaction partner of GATA1 is PU.1, crucial for regulating erythro-myeloid commitment. However, the nature and existence of the GATA1-PU.1 switch model are still a matter of discussion in the field. In our study, we do find co-expression of both GATA1 and PU.1 protein at the single-cell level. Importantly, PU.1 upregulation (at the mRNA and protein level), increased activity (ATAC-seq), and increased expression of target genes in *Gata1^mcherrry^* mutant cells strongly support the hypothesis that PU.1 and GATA1 are direct antagonists. Future studies will have to further elucidate the functions of GATA1 domains, in particular regarding the potential intramolecular interaction between the two transactivating domains, as well as their interaction with PU.1.

### Extrinsic factors influencing megakaryocyte hyperproliferation

*Gata1^mCherry^* blocks Mk differentiation exclusively in yolk sac-derived and not in intraembryonic-derived megakaryopoiesis. These ontogeny-specific cell fates could be governed by both intrinsic differences and distinct extrinsic cues.

For instance, it was proposed that the higher level of type 1 interferon in the adult bone marrow compared to the fetal liver inhibits Mk hyperproliferation, thereby contributing to the lack of Mk phenotype in adult *Gata1s* animals (Woo et al., 2013). Similarly, the fetal liver but not fetal bone marrow stroma support blast growth in co-cultures (Miyauchi & Kawaguchi, 2014).

Additionally, it is conceivable that EMP- and HSPC-derived Mks may compete for cytokines, as recently shown for the erythroid lineage (Soares-da-Silva et al., 2021). As EMP-derived Mks emerge and colonize the FL before HSPC-derived counterparts, accumulation of EMP-derived Mks in *Gata1^mcherry^* embryos may limit the availability of cytokines for HSPC-derived Mks and thus prevent them from hyperproliferating. In our study, female embryos, however, provide evidence that this is not the case. We took advantage of the fact that *Gata1* is encoded on the X-chromosome and is subjected to X-inactivation. Thus, female heterozygous embryos are chimeras containing both *Gata1^wild-type^*- and *Gata1^mCherry^*-expressing cells, that can be distinguished by mCherry expression. In female embryos, the hyperproliferation of *Gata1^mCherry^* cells did not perturb the differentiation of wild-type Mks. This indicates that *Gata1^mCherry^*-driven Mk hyperproliferation is not sufficient to inhibit megakaryopoiesis from other lineages co-existing simultaneously in the same niche.

### Intrinsic differences between yolk sac- and intraembryonic-derived hematopoiesis driving stage-specific phenotypes

Fetal Mks have been described to be intrinsically more proliferative than their adult counterparts (Slayton et al., 2005). Furthermore, in humans, hematopoietic progenitors from the fetal liver, cord blood, and adult BM respond differently when a *Gata1s* mutation is introduced *in vitro* (Gialesaki et al., 2018), suggesting an intrinsic difference between fetal and adult Mks. Indeed, Klusmann and colleagues showed that IGF2 signaling was required for *Gata1s*-induced hyperproliferation in fetal liver progenitor cells but that IGF2 had no effect on bone marrow-derived cells. They showed that *Gata1s* acted downstream of the IGF2 signaling cascade by failing to repress E2F leading to the overexpression of proliferation genes such as *Myc* (Klusmann et al., 2010).

In addition, the timespan from the emergence of the progenitor to the differentiation of the mature Mk differs between yolk sac- and the HSPC-derived Mks. While EMPs arise on day E8.5 and produce Mks 24-48 hours later (Palis et al., 1999; Tober et al., 2007; Iturri et al., 2021), there are at least 4 days (96 hours) between the emergence of an HSPC and their differentiation into Mks (Medvinsky et al., 1993; Müller et al., 1994). This could provide the cells more time to compensate for the lack of full-length Gata1 and to induce alternative mechanisms that prevent the hyperproliferation of Mks and the development of blast-like cells. This longer duration also indicates that in wild-type conditions, the signaling processes in the differentiation and maturation of HSPC-derived Mks may be fundamentally different from that of EMP-derived MKs, as we showed.

Finally, our data revealed that megakaryopoiesis is transcriptionally differently regulated and passes through distinct immunophenotypic populations at the cellular level depending on the stage and ontogeny. This is in line with the findings of a previous study (Cortegano et al., 2019) and could explain the stage-specific nature of *Gata1* mutants. Strikingly, those transcription factors *(Cebpb*, *Ikzf2*, *Pu.1,* and *Relb)* that showed stage-specific expression levels in the wild-type were also the transcription factors that had increased activity in mutant Mks and in the blast-like cells. This indicates that the different intrinsic properties of hematopoietic waves do indeed determine which phenotype a mutation will exert.

A signaling pathway required only for HSPC but not EMP emergence in mice is Notch signaling (Hadland et al., 2004; Robert-Moreno et al., 2008). Interestingly, Notch is regulated by NF-kB signaling in zebrafish hematopoiesis (Espín-Palazón & Traver, 2016), and Relb, one of the transcription factors particularly active at E16.5 is a member of the NF-kB family (Oeckinghaus & Ghosh, 2009). This suggests that NF-kB signaling might differently affect EMP- and HSPC-derived hematopoiesis. Furthermore, we found that a Gata2 binding site in a candidate cis-regulatory elements (cCRE) of *Relb* was differently accessible between mutant and wild-type cells in an ontogeny-specific manner. Interestingly, it has been proposed that Gata2 has ontogeny-specific functions and can compensate for lack of Gata1 (Fujiwara et al., 2004; Huang et al., 2009; Silvério-Alves et al., 2023).

Collectively, these studies indicate that lineage commitment and differentiation from distinct hematopoietic waves are controlled by unique molecular mechanisms and respond differently to genetic perturbations.

### *Gata1^mCherry^* causes the development of blast-like cells

*Gata1^mCherry^* not only causes the transient accumulation of immature Mks in yolk sac-derived lineages but also drives the development of blast-like cells. These blast-like cells are primed in the yolk sac and further mature in the fetal liver, characterized by the expression of CD244 as well as increased expression and activity of several myeloid and pluripotency transcription factors such as PU.1 (*Spi1*) and Helios (*Ikzf2*). CD244 is expressed by various immune cells such as natural killer (NK) cells as well as MPPs (Garni-Wagner et al., 1993; Kiel et al., 2005). We confirmed that blast-like cells are not NK cells (data not shown). This indicates that CD244 expression is here a marker of loss of commitment. The characterization of the cell surface receptor CD244 as a marker for at least a subset of Gata1 mutant-driven blast-like cells will allow future studies to investigate whether blast-like cells in *Gata1s* mouse models share similar transcriptomic and functional properties.

In line with human *Gata1s* expressing blasts that have the potential for eosinophils, mast cells, and megakaryocytes (Miyauchi et al., 2010; Maroz et al., 2014), CD244^+^ blast-like cells showed granulocyte, mast cell, and erythrocyte gene expression patterns. In *Gata1^mCherry^*, however, CD244^+^ blast-like cells only differentiated into CD41^low^ undifferentiated cells and erythrocytes and not into myeloid cells *in vitro*.

### Transcriptional induction of a blast-like cell fate

We observed that CEBPB was the most active transcription factor in *Gata1^mCherry^* Mks and CD244^+^ blast-like cells at E10.5. *Cebpb* is associated with myeloid and particularly granulocyte fate (Ness et al., 1993; Graf & Enver, 2009; Mancini et al., 2012; Cirovic et al., 2017). Later, at E12.5, the number of transcription factors with heightened activity in *Gata1^mCherry^* embryos increased, with Helios (*Ikzf2*, regulator of multipotent hematopoietic progenitors (Cova et al., 2021) and regulatory T cell development (Hahm et al., 1998; Thornton et al., 2010)) at the top. While these transcription factors are both important for lineage commitment and differentiation, they have also been shown to be involved in the maintenance of pluripotency (Cova et al., 2021) and the development of leukemia (Park et al., 2019; Yusenko et al., 2021; Cova et al., 2021; Klempnauer, 2022).

CEBPB has been reported to cooperate with P300 and MYB in leukemic cells (Yusenko et al., 2021; Klempnauer, 2022). In our study, *Myb* deficiency did not affect blast-like cell development. This indicates that in the context of *Gata1^mCherry^,* CEBPB regulates blast-like fate independently of MYB.

Helios has been reported to repress Mk fate in order to maintain pluripotency at the level of hematopoietic stem and progenitor cells in the adult bone marrow (Cova et al., 2021). This is in line with our finding that CD244^+^ blast-like cells which express high levels of *Ikzf2* have reduced megakaryocyte output in contrast to CD244^neg^ cells.

### Prenatal origin of pediatric leukemias in humans

As in mice, the human hematopoietic system is established in several sequential and overlapping waves originating from spatiotemporally distinct regions (Calvanese et al., 2022). This led to the hypothesis that childhood leukemias might originate from fetal progenitors (Cazzola et al., 2021; Camiolo, Mullen & Ottersbach, 2024).

One example of a pre-leukemic disorder that originates *in utero* is transient abnormal myelopoiesis (TAM). It is caused by mutations in the 2^nd^ exon of *Gata1* found in around 10 % of all children with Down Syndrome. These mutations lead to the expression of only the short isoform of *Gata1*, *Gata1s(short)* (Mundschau et al., 2003; Rainis et al., 2003). *Gata1s* causes hyperproliferation of megakaryoblasts in these children already *in utero* in the fetal liver (Smrcek et al., 2001; Zipursky, 2003). In most cases, TAM spontaneously resolves within the first 3 months after birth. Only if subsequent mutations occur, TAM further develops into acute megakaryoblastic leukemia (Labuhn et al., 2019; Sato et al., 2024). It has already been demonstrated through *in vitro* experiments that only human fetal liver, but not human cord blood or human bone marrow-derived Mks accumulate in the presence of GATA1s (Gialesaki et al., 2018). Unfortunately, the current lack of tools in human embryonic research to ascribe ontogeny to fetal cells has hampered the identification of the exact progenitor origin of the studied fetal cells.

Additionally, studies performed in mice suggest that also pediatric acute lymphoid leukemias (ALLs) might originate from distinct cellular origins than adult ALL.

For example, Sinha and colleagues demonstrated that MLL-ENL translocation causes a more aggressive disease when induced at E12.5 than in the bone marrow (Sinha et al., 2020). Furthermore, another study also showed that the expression of the MLL-AF4 fusion gene causes a stronger effect between E12.5 and E14.5 than at later stages (Barrett et al., 2016). This effect was associated with lymphoid-primed multipotent progenitors (LMPPs) (Böiers et al., 2013) which are a distinct population of HSPCs that most likely rely on unique molecular mechanisms. Altogether, these studies support our finding that ontogeny determines the effect a mutation exerts.

As the landscape of the different hematopoietic progenitor waves and their intrinsic particularities are being refined in mice and humans, our understanding of the origins and molecular events of the onset of infant and pediatric blood disorders will increase.

### Limitations of the study

This study identified for the first time the stage-specific perturbations induced by a fusion GATA1^mCherry^ protein that reproduced the effects of another known *Gata1* mutant, *Gata1s*. It would be important to confirm whether CD244^+^ blast-like cells which we discovered in *Gata1^mCherry^* embryos are also present in the fetal livers of *Gata1s* mice and in human DS-TAM patients. Furthermore, our single-cell data does not allow us to pinpoint the exact cellular stage within the trajectory from multipotent progenitor to Mk at which *Gata1^mCherry^*exerts its effect and perturbs megakaryopoiesis at the transcriptional level.

There is increasing evidence that there is an additional hematopoietic progenitor wave emerging after EMPs and before HSPCs called embryonic multipotent progenitors (eMPPs) (Patel et al., 2022; Yokomizo et al., 2022; Barone et al., 2024; Soares-da-Silva et al., 2025). Since the contribution of this wave to megakaryocyte production is yet to be fully uncovered and as it remains challenging to specifically label this wave without simultaneously labeling EMPs or HSPCs, we decided to here focus only on EMPs (*Cdh5^CreERT2^* pulsed at E7.5 or *Csf1r^MeriCreMer^*pulsed at E8.5) and HSPCs (*Cdh5^CreERT2^* pulsed at E10.5). Importantly, pulse labeling in *Csf1r^MeriCreMer^* pulsed at E8.5 does not label this eMPP population. It will, however, be interesting to test in the future whether eMPP-derived Mks are affected or not by *Gata1^mCherry^*.

## Conclusion

In summary, we provide evidence that the ontogeny or cellular origin determines which phenotype or disease a mutant/mutation will exert. This has the potential to help further elucidate the origins of infant and pediatric diseases such as leukemias.

## Acknowledgments

We thank all team members for their support and constructive discussion. We thank Timm Schroeder for kindly sharing the Gata1^mCherry^ mice and Peggy Kirstetter (IGBMC, Illkrich, France) for insight into their single-cell methocult protocol. We acknowledge the flow cytometry platform (Sebastien Megharba, Sandrine Schmutz, and Sophie Novault) and single-cell biomarkers platform (Carolina Moraes Cabé, Valérie Seffer) of the CB UTechS at Institut Pasteur for support in conducting this study. We are grateful to all personnel of the Institut Pasteur center for Animal resources and research for their invaluable work. We thank Yann Loe Mie with support in initial analysis of the sequencing data. We thank Azimdine Habib and Marc Monot from the Biomics Platform, C2RT, Institut Pasteur, Paris, France, supported by France Génomique (ANR-10-INBS-09) and IBISA. This work was supported by recurrent funding from Institut Pasteur, CNRS, and Revive (Investissement d’Avenir; ANR-10-LABX-73), by an ERC investigator award (2016-StG-715320) from the European Research Council and by Impulscience® from Fondation Bettencourt Schueller to E.G.P. A.S. was funded by a fellowship from FRM (FDT202304016740).

## Data availability

Paired single-cell RNA and ATACseq data will be made available on a public repository.

## Code availability

The code of our single-cell RNA and ATACseq data will be made available.

## Author contributions

Conceptualization: A.S and E.G.P.

Methodology and data collection: Flow cytometry and animal breeding: A.S., L.F., Y.L., and P.D.

scMultiome-seq data analysis: S.F., A.S., and E.G.P.

Writing - Original Draft: A.S.; Writing - Review & Editing: A.S., S.F., L.F., and E.G.P.

Funding Acquisition: E.G.P.

Supervision: E.G.P.

