## Supplement for "Ontogeny drives stage-specific effects of a Gata1 mutation"

### Methods

#### Mice

All mice used in this study have been previously described. Experimental procedures, housing, and husbandry were in compliance with the regulatory guidelines of the Institut Pasteur Committee for Ethics and Animal Experimentation (CETEA, dap160091 and dap190119). Strains included *Csf1r*$^{iCre}$ (FVB background, MGI:4429470) (Deng et al., 2010), *Csf1r*$^{MeriCreMer}$ (FVB background, MGI:J:186831) (Qian et al., 2011), *Cdh5*$^{CreERT2}$ (C57Bl/6 background, MGI:3848982) (Sörensen et al., 2009), *PGK*$^{Cre}$ transgenic mice (C57Bl/6 background, MGI:2178050) (Lallemand et al., 1998), *Gata1*$^{mCherry}$ (kind gift from Timm Schroeder, ETH Zürich, C57Bl/6 background) (Hoppe et al., 2016), *c-Myb* mutant (Mucenski et al., 1991), *Rosa26*$^{(e)YFP}$ (C57Bl/6 background, MGI:J:80963) (Srinivas et al., 2001) and *Rosa26*$^{Fucci2aR}$ (C57Bl/6 background, MGI:5645798) (Mort et al., 2014). Timed matings were performed and the date of the vaginal plug was considered E0.5. Embryonic stages were validated using somite counting and morphological landmarks.

#### Genotyping

PCR genotyping of *Csf1r*$^{iCre}$ (Deng et al., 2010), *Csf1r*$^{MeriCreMer}$ (Qian et al., 2011), *Cdh5*$^{CreERT2}$ (Sörensen et al., 2009), *Gata*$^{mCherry}$ (Hoppe et al., 2016), *c-Myb* (Mucenski et al., 1991)*, and R26*$^{Fucci2aR}$ (Mort et al., 2014)*, R26*$^{YFP}$ (Srinivas et al., 2001) embryos, and mice was performed according to protocols described earlier and are available upon request.

#### 4-OHT preparation and injection

4-hydroxytamoxifen (4-OHT) (Sigma, H7904-25MG) was dissolved in equal parts of ethanol and Kolliphor (Sigma C5135-500G) using sonication. 10 mg/mL stocks of progesterone (P3972-5G) were prepared by resuspending in ethanol and sunflower oil (Sigma S5007-250ML) and co-injected with OHT to reduce the risk of abortion. For pulse-labeling using the *Csf1r*$^{MeriCreMer}$ strain, females were weighed on day 8 of pregnancy and injected with 75 µg/g (body weight) OHT and 37.5 µg/g (body weight) progesterone at 1 pm. For pulse-labeling using the *Cdh5*$^{CreERT2}$ strain, females were weighed on day 7 or 10 of pregnancy and injected with 50 µg/g (body weight) OHT and 25 µg/g (body weight) progesterone at 1 pm.

**Flow cytometry**

Pregnant mice were killed by cervical dislocation and embryos were dissected in cold PBS. Fetal peripheral blood was collected in 2 mM EDTA by severing the vitelline and umbilical vessels after removing the placenta and extraembryonic membranes. Fetal organs were enzymatically dissociated in digestion buffer composed of PBS with 1 mg/mL collagenase D (Sigma 11088882001), 100 U/mL DNaseI (DN25-100mg), and 3% Fetal Bovine Serum for 30 minutes at 37 °C. Fetal organs were passed through 100 µm strainers by mashing with the piston of a 2 mL syringe and then collected in ice-cold filtered FACS Buffer (0.5% BSA and 2mM EDTA in PBS). Blocking was performed with 5% FBS and 1:20 mouse IgG (Interchim 015-000-003) in FACS Buffer followed by 30 minutes of antibody staining (antibodies listed below). Cells were washed and incubated with fluorescently-conjugated streptavidin for 20 minutes when applicable. Stained cells were passed on the Cytoflex LX (Beckman Coulter) or BD Symphony A5 (BD Biosciences). To quantify cells, the following formula was used : (# of cells acquired) x (volume of resuspended cells after staining and washing/volume of cells acquired) x (volume of cell suspension in blocking buffer before staining/volume of cells plated for staining). Results were analyzed and plots were generated using FlowJo™ v10.10.0 Software (BD Life Sciences).

| **Antigen** | **Clone** |
|---|---|
| CD16/32 | 2.4G2 |
| CD45 | 30-F11 |
| CD41 | MWReg30 |
| F4/80 | BM8 |
| Ly6G | 1A8 |
| Ly6C | HK1.4 |
| Sca1 | D7 |
| CD34 | RAM34 |
| Ter119 | TER-119 |
| CD71 | C2 |
| CD19 | 1D3 |
| CD3e | 145-2C11 |
| Gr1 | RB6-8C5 |
| NK1.1 | PK136 |
| Kit | 2B8 |
| CD11b | M1/70 |
| CD150 | mShad150 |
| CD48 | HM48-1 |
| CD244.2 | REAL493 |
| CD42d | 1C2 |
| Ki67 | B56 |
| PU.1 | E.388.3 |
| Helios | 22F6 |

**Intracellular staining for flow cytometry**

Cells were prepared for flow cytometry as described above. After cell surface staining, cells were fixed and permeabilized for 1 h using the FoxP3 kit (Invitrogen 00-5523-00) by following the manufacturer's instructions. Afterward, cells were washed and stained for 1 h with primary antibody in permeabilization buffer at room temperature on a shaker (1:100 rabbit anti-PU.1 (Invitrogen MA5-15064, clone E.388.3), 1:100 Ki67-AF647 (BD558615, clone B56), or 1:100 Helios-APC (Biolegend 137222, clone 22F6)) and washed again. In the case of the PU.1 staining, secondary antibody staining was performed for 1 h in permeabilization buffer under the same conditions as the primary antibody (anti-rabbit AF647 (Invitrogen 10123672), 1:100).

**Hoechst staining for polyploidy assay**

Cells were prepared for flow cytometry as described above. Cells were then incubated with FACS buffer containing 1:100 Hoechst 33342 (Miltenyi 130-111-569) in a shaking incubator at 37 °C with 100 rpm for 40 min. Single cell suspensions were washed and acquired on the flow cytometer.

**Immunofluorescence of yolk sac whole mounts**

Yolk sacs were fixed in 10 % Formalin solution containing 4 % formaldehyde (Sigma) for 2 h at 4 °C and washed 3 times in 1x PBS at room temperature. Yolk sacs were then permeabilized and blocked in 10 % Normal Goat Serum in PBS-Triton 0.5% (PBS-T) for 2 h at room temperature and stained in blocking buffer containing rat anti-CD41-PE (1:100; clone MWREg 30 Biolegend) overnight at 4 °C. Finally, yolk sacs were washed 3 times for 10 min in PBS-T at room temperature and mounted in ProLong$^{TM}$ Gold antifade (Life technologies, P36930). Z-stacks were acquired on a *Leica SP8* confocal microscope with a 40x/1.30NA objective (immersion oil). Maximum projections were generated in *Fiji* (Schindelin et al., 2012) and PE exposure was adapted for better visibility of the signal.

**Immunofluorescence of thick fetal liver sections**

Fresh 100 µm thick E12.5 or E16.5 fetal liver sections were prepared as previously described (Peixoto et al., 2024). Dissected fetal livers (FL) were fixed in 5 % Formalin solution containing 2 % formaldehyde (Sigma, HT5014) in PBS at 4 °C overnight and washed 3 times in PBS for 15 min each. 100 µm thick sections were cut in PBS using a vibratome (Leica, VT 1200S). For E16.5 samples, sections were permeabilized and blocked in 10% Normal Goat Serum in PBS-Triton 0.5% (PBS-T) for 2 h at room temperature, washed 3x 20 min in 0.1 % PBS-T, incubated with chicken anti-GFP (1:100; Abcam ab13970) in blocking solution overnight at 4 °C and finally washed again 3x 20 min in 0.1 % PBS-T. Secondary staining was performed by incubation with anti-chicken AF488 (1:500; Invitrogen #10286672) and rat anti-CD41-PE (1:100; clone MWREg 30 Biolegend) in 0.1 % PBS-T for 2 h at room temperature in the dark. FL sections were washed 3x 20 min in 0.1 % PBS-T and cleared in RapiClear (1.52; Sunjin Lab RC152001) overnight. For E12.5 samples, the same protocol was followed but staining was only performed using rat anti-CD41-PE. All sections were mounted in RapiClear and z-stacks were acquired as described above.

**Single-cell liquid culture**

Fetal livers were dissected and blocked as described above. Lineage cells (Ter119$^+$ CD19$^+$ CD3e$^+$ CD4$^+$ CD8$^+$ NK1.1$^+$ F4/80$^+$ Gr1$^+$) were depleted using magnetic anti-biotin Microbeads (1:5) (Miltenyi Biotec 130-090-485) and MS columns (Miltenyi Biotec 130-042-201). Single cells were sorted using a FACSAria III (Diva software) into flat-bottom 96-well plates containing pre-warmed and equilibrated differentiation medium (0.1% β-mercaptoethanol, 1X Penicillin/Streptomycin, 10% FBS, 1:125 SCF, 5ng/µL GM-CSF, 5ng/µL M-CSF, 2ng/µL EPO and 5ng/µL TPO in Opti-MEM with Glutamax). SCF was supplied from myeloma cell line supernatant. Cells were grown at 37°C with 5% $CO_2$ for 7 days. Colonies were manually scored to detect small megakaryocyte colonies before collecting them by scratching with a pipette tip for flow cytometry analysis. The overall cloning efficiency was defined as the number of sorted single cells that give rise to colonies. The Mk cloning efficiency was defined as the number of sorted single cells that give rise to Mk containing colonies. The colony output was calculated by dividing the number of colonies of one type, i.e. erythrocytes, by the number of all colonies grown.

**Single-cell MethoCult culture**

Single cells were sorted into 96-well plates containing 50 µL of MethoCult™ GF M3434 (Stemcell Technologies, 03434) as described above. Cells were grown at 37 °C with 5 % $CO_2$ for 10 days. Colonies were manually scored to detect small colonies before washing them twice with 1x PBS and staining them for flow cytometry analysis.

#### Single-cell multiome experiment

##### *Library preparation*

Cells were sorted using a Symphony S6 (BD Biosciences) with an 85 µL m nozzle into 350 µL PBS containing 5 % FBS. Paired scRNA-and scATAC-seq libraries were prepared using the 10x Genomics Single-Cell Multiome ATAC + Gene Expression kit according to the manufacturer's instructions. The initial lysis step was performed for 4 min. ATAC sample index PCR was run with 8 cycles, the cDNA amplification step with 9 cycles, and the GEX sample index PCR with 11 cycles.

Libraries were sequenced with paired-end 100-bp reads on a NextSeq 2000 (Illumina) to a target depth of 25,000 read pairs per nucleus for the ATAC library and 25,000 read pairs per nucleus for the GEX library.

##### *Analysis and annotation of the 10x multiome dataset*

Sequencing reads from the six multiome samples were processed using 10X's Cell Ranger ARC pipeline (mm10-2020A genome) to produce count matrices, which were then further processed using the R packages Seurat (Hao et al., 2021) and Signac (Stuart et al., 2021). To combine datasets from the same time points, we computed time point-specific consensus peak sets by merging overlapping peaks from the two conditions, keeping all peaks smaller than 10 kbp. We then re-quantified the count matrices using Signac's FeatureMatrix function. Low-quality cells were filtered out according to the following criteria: >100,000 RNA counts, >100,000 ATAC counts, <30% reads in peaks, >5% of ATAC reads mapping to ENCODE blacklisted regions (Amemiya et al., 2019), >2 nucleosome signal, <2 TSS enrichment, doublet score >0.75 (assessed using DoubletFinder (McGinnis et al., 2019) for RNA and scDblFinder for ATAC (Germain et al., 2022)). We used the following sample-specific minimal thresholds: >1,500 genes for E10.5-wt, E10.5-*Gata1*$^{mCherry}$, E12.5-wt, >1,000 genes for E12.5- *Gata1*$^{mCherry}$ and E16.5- *Gata1*$^{mCherry}$, >500 genes for E16.5 wt, >5,000 ATAC counts for E12.5- *Gata1*$^{mCherry}$, >2,000 ATAC counts for all other datasets.

We ran two analyses: an unintegrated analysis of each time point that highlights fine biological distinctions between wild-type and *Gata1*$^{mCherry}$ mutants and an integrated analysis of the RNA modality to harmonize annotations and visualize all data in a single embedding. For each time point, we combined the RNA and ATAC count matrices (without integration). For the RNA modality, we computed log-normalized counts (counts per 10K), variable features (top 2000 genes, excluding mitochondrial genes and gene models), scaling, PCA, UMAP, SNN, and Louvain clustering. For the ATAC modality, we computed the TF-IDF normalization, SVD (removing the first component), UMAP, SNN, and Louvain clustering. We computed Seurat's WNN analysis (Hao et al., 2021) to obtain a multiome UMAP and clustering. We chose to use the unintegrated RNA analysis to display fine differences at each time point as the RNA modality produced the finest clustering and visualization results, while systematic differences between conditions (technical effects, sex) were negligible compared to biological effects at all time points (good mixing of samples around the MPP compartment, perfect mixing of males and females). To obtain harmonized annotations and visualizations for all time points and conditions, we combined and re-processed the RNA modality of all six samples. We applied Harmony integration (Korsunsky et al., 2019) to obtain a joint PCA-like embedding, then computed UMAP, SNN, and Louvain clustering. We manually annotated the resulting clusters using known marker genes.

***Analysis and annotation of the 10x multiome dataset***

To mitigate batch effects (ambient content, male vs female differences), we used generalized linear models with the following formula:

$$g(Y) = \alpha.cell\_type + \beta.sample + \gamma.cell\_type.sample + \delta.log\_counts$$

where $Y$ are expression values (normalized or counts depending on the modality), $cell\_type$ is an indicator function (one-vs-all setting, 1 for the cell type of interest, 0 for cells from other types), $log\_counts$ is the per-cell log number of counts (used when the model is fit on counts). Coefficient $\alpha$ captures cell-type related effects, $\beta$ captures systematic biological and technical variablity, $\gamma$ captures cell-type specific conditional effects. In the paper, we focused on the interpretation of $\alpha$ (cell-type effects) and $\gamma$ (within cell-type conditional effects). GLM computations and p-value estimations were performed independently at each time point using the fastglm package (Huling et al., 2022), then FDR-corrected (Benjamini & Hochberg, 1995). For the RNA modality, we applied a plain linear model on ranked normalized expression. For the ATAC modality, we used a Poisson model on counts, keeping only peaks detected in >100 cells.

##### *Gene regulatory networks*

To compute transcription factor activity at the single-cell level in the ATAC modality, we ran ChromVAR (Schep et al., 2017) using Signac's RunChromVAR function on the JASPAR 2024 motif set (all mammals, (Rauluseviciute et al., 2024)). To compare differential activity at the cluster level, we fit two sets of linear models: across cell types (pooled conditions, one cell type against all others) and across conditions (one model per cell type). For each model, we used the log number of fragments as a covariable to adjust for sequencing depth.

To compute gene regulatory networks (GRNs), we applied Pando (Fleck et al., 2023) at each time point, pooling conditions. Pando finds triplets of transcription factors, candidate cis-regulatory elements (cCREs), and target genes using a generalized linear model at the single-cell level (independently of clusters) that directly combines RNA and ATAC information. To focus on regulations on the megakaryocyte branch, we subsetted the data to MEPs, megakaryocytes and megakaryocyte progenitors. To obtain robust results, we aggregated cells into metacells using the Louvain algorithm at a resolution of 20 (~50 cells per metacell) and ran Pando with the XGB algorithm on all genes detected in >1% cells. We used the JASPAR 2024 motif set to initiate the GRN. For each TF, the XGB algorithm finds a regulon consisting of up to 50 cCREs and target genes. We then estimated the activity of each regulon at the single-cell level for each modality: on the RNA side, we computed an AUCell-like score (Aibar et al., 2017) using a standard Area under the ROC Curve (scoring the rank of target genes among all genes); on the ATAC side, we computed the ChromVAR score for the top cCREs of each TF. Finally, we computed differential regulon activity across cell types and conditions using linear models (see ChromVAR differential activity), using the log number of fragments as a covariable for the ATAC, without covariable for the RNA.

##### *Pseudotime and co-expression analysis*

We computed pseudotime values using Palantir (Setty et al., 2019) on the joint Harmony embedding, with multipotent progenitors as the starting point and all other clusters except MEP as endpoints. To focus on expression differences along the megakaryocyte branch, we extracted Mk I and Mk II, then re-computed markers for each cell type and condition combination at each time point using MetaMarkers (Fischer & Gillis, 2021). We extracted all genes with both high cell-type variability (AUROC>0.6, FDR<$10^{-5}$ in any cell-type by condition combination) and high conditional variability (log2FC>0.5, FDR<0.01 specifically for the WT vs mutant Mk I comparison). We then grouped these genes by co-expression along the MEP to megakaryocyte differentiation path: we computed the Pearson correlation of expression values in MEP, Mk I, and Mk II at each time point, rank-normalized the resulting co-expression matrices, then averaged the matrices across the 3 timepoints. We obtained co-expression modules by applying the dynamic tree cut algorithm (Langfelder et al., 2008) to the hierarchical clustering with average linkage of the average coexpression matrix. Finally, we computed the average expression of each module in all cells using the MetaMarkers score_cells function. To account for sequencing depth differences, the sum of expression across modules was normalized to 1 for each cell, then module expression scores were z-scored across cells. For visualization purposes, module scores were smoothed along pseudotime using a window size of 0.2. We performed gene enrichment analysis of the genes of module 1+2, 3 and 4+5 using Enrichr (Chen et al., 2013; Kuleshov et al., 2016; Xie et al., 2021).

**Statistics and Graphs**

Statistical analysis was performed with GraphPad Prism v10.1.1. Figures were created with GraphPad Prism v10.1, Biorender.com and Adobe Illustrator v27.9.1.

### Methods – References


Aibar, S., González-Blas, C. B., Moerman, T., Huynh-Thu, V. A., Imrichova, H., Hulselmans, G., Rambow, F., Marine, J. C., Geurts, P., Aerts, J., Van Den Oord, J., Atak, Z. K., Wouters, J., & Aerts, S. (2017). SCENIC: Single-cell regulatory network inference and clustering. *Nature Methods*, *14*(11), 1083–1086. https://doi.org/10.1038/nmeth.4463

Amemiya, H. M., Kundaje, A., & Boyle, A. P. (2019). The ENCODE Blacklist: Identification of Problematic Regions of the Genome. *Scientific Reports*, *9*(1), 9354. https://doi.org/10.1038/s41598-019-45839-z

Benjamini, Y., & Hochberg, Y. (1995). Controlling the False Discovery Rate: A Practical and Powerful Approach to Multiple Testing. *Journal of the Royal Statistical Society: Series B (Methodological)*, *57*(1), 289–300. https://doi.org/10.1111/j.2517-6161.1995.tb02031.x

Chen, E. Y., Tan, C. M., Kou, Y., Duan, Q., Wang, Z., Meirelles, G., Clark, N. R., & Ma'ayan, A. (2013). Enrichr: Interactive and collaborative HTML5 gene list enrichment analysis tool. *BMC Bioinformatics*, *14*(1), 128. https://doi.org/10.1186/1471-2105-14-128

Deng, L., Zhou, J.-F., Sellers, R. S., Li, J.-F., Nguyen, A. V., Wang, Y., Orlofsky, A., Liu, Q., Hume, D. A., Pollard, J. W., Augenlicht, L., & Lin, E. Y. (2010). A Novel Mouse Model of Inflammatory Bowel Disease Links Mammalian Target of Rapamycin-Dependent Hyperproliferation of Colonic Epithelium to Inflammation-Associated Tumorigenesis. *The American Journal of Pathology*, *176*(2), 952–967. https://doi.org/10.2353/ajpath.2010.090622

Fischer, S., & Gillis, J. (2021). How many markers are needed to robustly determine a cell's type? *iScience*, *24*(11), 103292. https://doi.org/10.1016/j.isci.2021.103292

Fleck, J. S., Jansen, S. M. J., Wollny, D., Zenk, F., Seimiya, M., Jain, A., Okamoto, R., Santel, M., He, Z., Camp, J. G., & Treutlein, B. (2023). Inferring and perturbing cell fate regulomes in human brain organoids. *Nature*, *621*(7978), 365–372. https://doi.org/10.1038/s41586-022-05279-8

Germain, P.-L., Lun, A., Meixide, C. G., Macnair, W., & Robinson, M. D. (2022). *Doublet identification in single-cell sequencing data using* scDblFinder (10:979). F1000Research. https://doi.org/10.12688/f1000research.73600.2

Hao, Y., Hao, S., Andersen-Nissen, E., Mauck, W. M., Zheng, S., Butler, A., Lee, M. J., Wilk, A. J., Darby, C., Zager, M., Hoffman, P., Stoeckius, M., Papalexi, E., Mimitou, E. P., Jain, J., Srivastava, A., Stuart, T., Fleming, L. M., Yeung, B., … Satija, R. (2021). Integrated analysis of multimodal single-cell data. *Cell*, *184*(13), 3573-3587.e29. https://doi.org/10.1016/j.cell.2021.04.048

Hoppe, P. S., Schwarzfischer, M., Loeffler, D., Kokkaliaris, K. D., Hilsenbeck, O., Moritz, N., Endele, M., Filipczyk, A., Gambardella, A., Ahmed, N., Etzrodt, M., Coutu, D. L., Rieger, M. A., Marr, C., Strasser, M. K., Schauberger, B., Burtscher, I., Ermakova, O., Bürger, A., … Schroeder, T. (2016). Early myeloid lineage choice is not initiated by random PU.1 to GATA1 protein ratios. *Nature*, *535*(7611), 299–302. https://doi.org/10.1038/nature18320

Huling, J., Bates, D., Eddelbuettel, D., Francois, R., & Qiu, Y. (2022). *fastglm: Fast and Stable Fitting of Generalized Linear Models using "RcppEigen"* (0.0.3) [Computer software]. https://cran.r-project.org/web/packages/fastglm/index.html

Korsunsky, I., Millard, N., Fan, J., Slowikowski, K., Zhang, F., Wei, K., Baglaenko, Y., Brenner, M., Loh, P., & Raychaudhuri, S. (2019). Fast, sensitive and accurate integration of single-cell data with Harmony. *Nature Methods*, *16*(12), 1289–1296. https://doi.org/10.1038/s41592-019-0619-0

Kuleshov, M. V., Jones, M. R., Rouillard, A. D., Fernandez, N. F., Duan, Q., Wang, Z., Koplev, S., Jenkins, S. L., Jagodnik, K. M., Lachmann, A., McDermott, M. G., Monteiro, C. D., Gundersen, G. W., & Ma'ayan, A. (2016). Enrichr: A comprehensive gene set enrichment analysis web server 2016 update. *Nucleic Acids Research*, *44*(W1), W90–W97. https://doi.org/10.1093/nar/gkw377

Lallemand, Y., Luria, V., Haffner-Krausz, R., & Lonai, P. (1998). Maternally expressed PGK-Cre transgene as a tool for early and uniform activation of the Cre site-specific recombinase. *Transgenic Research*, *7*(2), 105–112. https://doi.org/10.1023/a:1008868325009

Langfelder, P., Zhang, B., & Horvath, S. (2008). Defining clusters from a hierarchical cluster tree: The Dynamic Tree Cut package for R. *Bioinformatics*, *24*(5), 719–720. https://doi.org/10.1093/bioinformatics/btm563

McGinnis, C. S., Murrow, L. M., & Gartner, Z. J. (2019). DoubletFinder: Doublet Detection in Single-Cell RNA Sequencing Data Using Artificial Nearest Neighbors. *Cell Systems*, *8*(4), 329-337.e4. https://doi.org/10.1016/j.cels.2019.03.003

Mort, R. L., Ford, M. J., Sakaue-Sawano, A., Lindstrom, N. O., Casadio, A., Douglas, A. T., Keighren, M. A., Hohenstein, P., Miyawaki, A., & Jackson, I. J. (2014). Fucci2a: A bicistronic cell cycle reporter that allows Cre mediated tissue specific expression in mice. *Cell Cycle (Georgetown, Tex.)*, *13*(17), 2681–2696. https://doi.org/10.4161/15384101.2015.945381

Mucenski, M. L., McLain, K., Kier, A. B., Swerdlow, S. H., Schreiner, C. M., Miller, T. A., Pietryga, D. W., Scott, W. J., & Potter, S. S. (1991). A functional c-myb gene is required for normal murine fetal hepatic hematopoiesis. *Cell*, *65*(4), 677–689. https://doi.org/10.1016/0092-8674(91)90099-K

Peixoto, M. M., Soares-da-Silva, F., Bonnet, V., Ronteix, G., Santos, R. F., Mailhe, M.-P., Feng, X., Pereira, J. P., Azzoni, E., Anselmi, G., Bruijn, M. de, Baroud, C. N., Pinto-do-Ó, P., & Cumano, A. (2024). *Spatiotemporal dynamics of cytokines expression dictate fetal liver hematopoiesis* (p. 2023.08.24.554612). bioRxiv. https://doi.org/10.1101/2023.08.24.554612

Qian, B. Z., Li, J., Zhang, H., Kitamura, T., Zhang, J., Campion, L. R., Kaiser, E. A., Snyder, L. A., & Pollard, J. W. (2011). CCL2 recruits inflammatory monocytes to facilitate breast-tumour metastasis. *Nature*, *475*(7355), 222–225. https://doi.org/10.1038/nature10138

Rauluseviciute, I., Riudavets-Puig, R., Blanc-Mathieu, R., Castro-Mondragon, J. A., Ferenc, K., Kumar, V., Lemma, R. B., Lucas, J., Chèneby, J., Baranasic, D., Khan, A., Fornes, O., Gundersen, S., Johansen, M., Hovig, E., Lenhard, B., Sandelin, A., Wasserman, W. W., Parcy, F., & Mathelier, A. (2024). JASPAR 2024: 20th anniversary of the open-access database of transcription factor binding profiles. *Nucleic Acids Research*, *52*(D1), D174–D182. https://doi.org/10.1093/nar/gkad1059

Schep, A. N., Wu, B., Buenrostro, J. D., & Greenleaf, W. J. (2017). chromVAR: Inferring transcription-factor-associated accessibility from single-cell epigenomic data. *Nature Methods*, *14*(10), 975–978. https://doi.org/10.1038/nmeth.4401

Schindelin, J., Arganda-Carreras, I., Frise, E., Kaynig, V., Longair, M., Pietzsch, T., Preibisch, S., Rueden, C., Saalfeld, S., Schmid, B., Tinevez, J. Y., White, D. J., Hartenstein, V., Eliceiri, K., Tomancak, P., & Cardona, A. (2012). Fiji: An open-source platform for biological-image analysis. *Nature Methods*, *9*(7), 676–682. https://doi.org/10.1038/nmeth.2019

Setty, M., Kiseliovas, V., Levine, J., Gayoso, A., Mazutis, L., & Pe'er, D. (2019). Characterization of cell fate probabilities in single-cell data with Palantir. *Nature Biotechnology*, *37*(4), 451–460. https://doi.org/10.1038/s41587-019-0068-4

Sörensen, I., Adams, R. H., & Gossler, A. (2009). DLL1-mediated Notch activation regulates endothelial identity in mouse fetal arteries. *Blood*, *113*(22), 5680–5688. https://doi.org/10.1182/blood-2008-08-174508

Srinivas, S., Watanabe, T., Lin, C. S., William, C. M., Tanabe, Y., Jessell, T. M., & Costantini, F. (2001). Cre reporter strains produced by targeted insertion of EYFP and ECFP into the ROSA26 locus. *BMC Developmental Biology*, *1*, 1–8. https://doi.org/10.1186/1471-213X-1-4.

Stuart, T., Srivastava, A., Madad, S., Lareau, C. A., & Satija, R. (2021). Single-cell chromatin state analysis with Signac. *Nature Methods*, *18*(11), 1333–1341. https://doi.org/10.1038/s41592-021-01282-5

Xie, Z., Bailey, A., Kuleshov, M. V., Clarke, D. J. B., Evangelista, J. E., Jenkins, S. L., Lachmann, A., Wojciechowicz, M. L., Kropiwnicki, E., Jagodnik, K. M., Jeon, M., & Ma'ayan, A. (2021). Gene Set Knowledge Discovery with Enrichr. *Current Protocols*, *1*(3), e90. https://doi.org/10.1002/cpz1.90

### Supplemental Data

#### Supplemental Data Figure 1

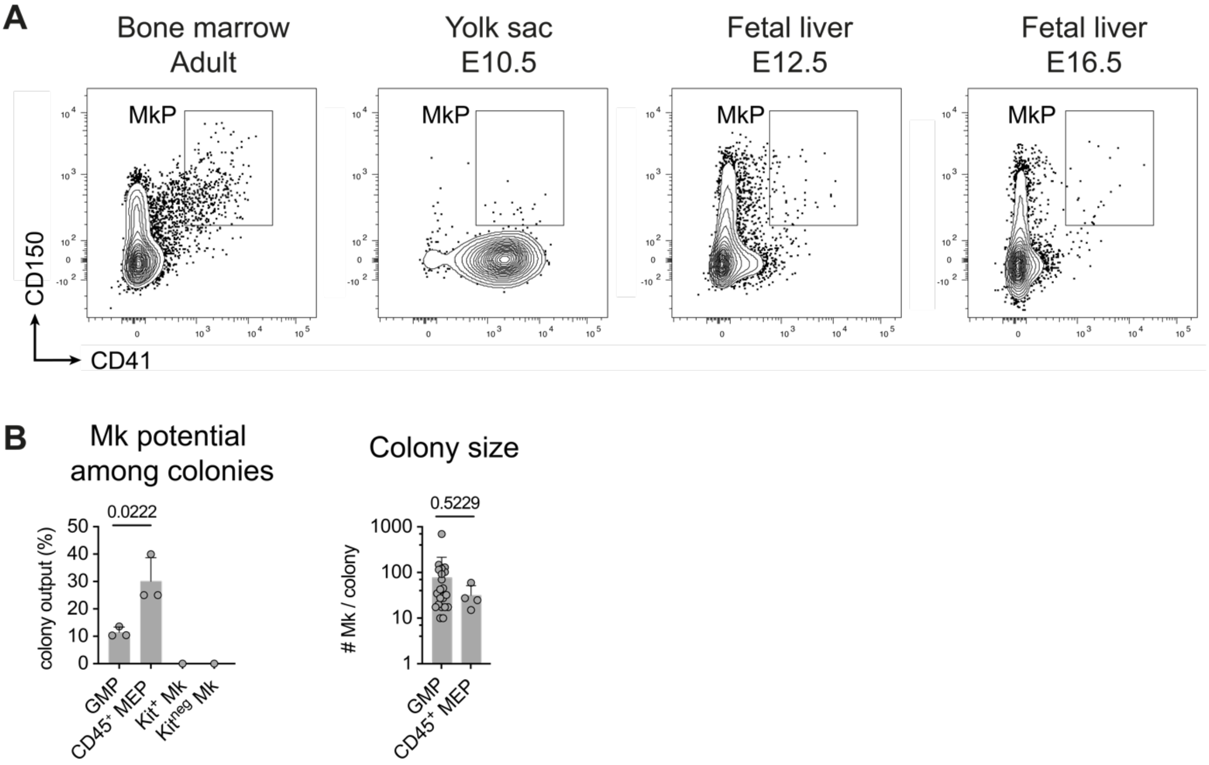


**Supplemental Data Figure 1, to Figure 1.** (**A**) Dot plot of $Lin^{neg}$ $Kit^{high}$ $Sca\text{-}1^{neg}$ cells in the adult bone marrow, E10.5 yolk sac, E12.5 fetal liver, and E16.5 fetal liver. Gate corresponds to $CD150^{+}$ $CD41^{+}$ Megakaryocyte progenitors (MkPs). (**B**) Mk potential among colonies (Mk cloning efficiency) (left) and colony size (right) grown from E12.5 fetal liver-derived immunophenotypic granulocyte-monocyte progenitors (GMPs, $Lin^{neg}$ $Kit^{high}$ $Sca\text{-}1^{neg}$ $CD16/32^{high}$ $CD34^{+}$), CD45+ megakaryocyte-erythrocyte progenitors (MEPs, $Lin^{neg}$ $Kit^{high}$ $Sca\text{-}1^{neg}$ $CD16/32^{neg}$ $CD34^{neg}$), $Kit^{+}$ $CD41^{+}$ and $Kit^{neg}$ $CD41^{+}$ Mks from 3 independent litters. Šidak multiple comparisons. Data are represented as mean ± SD.

**Supplemental Data Figure 2**

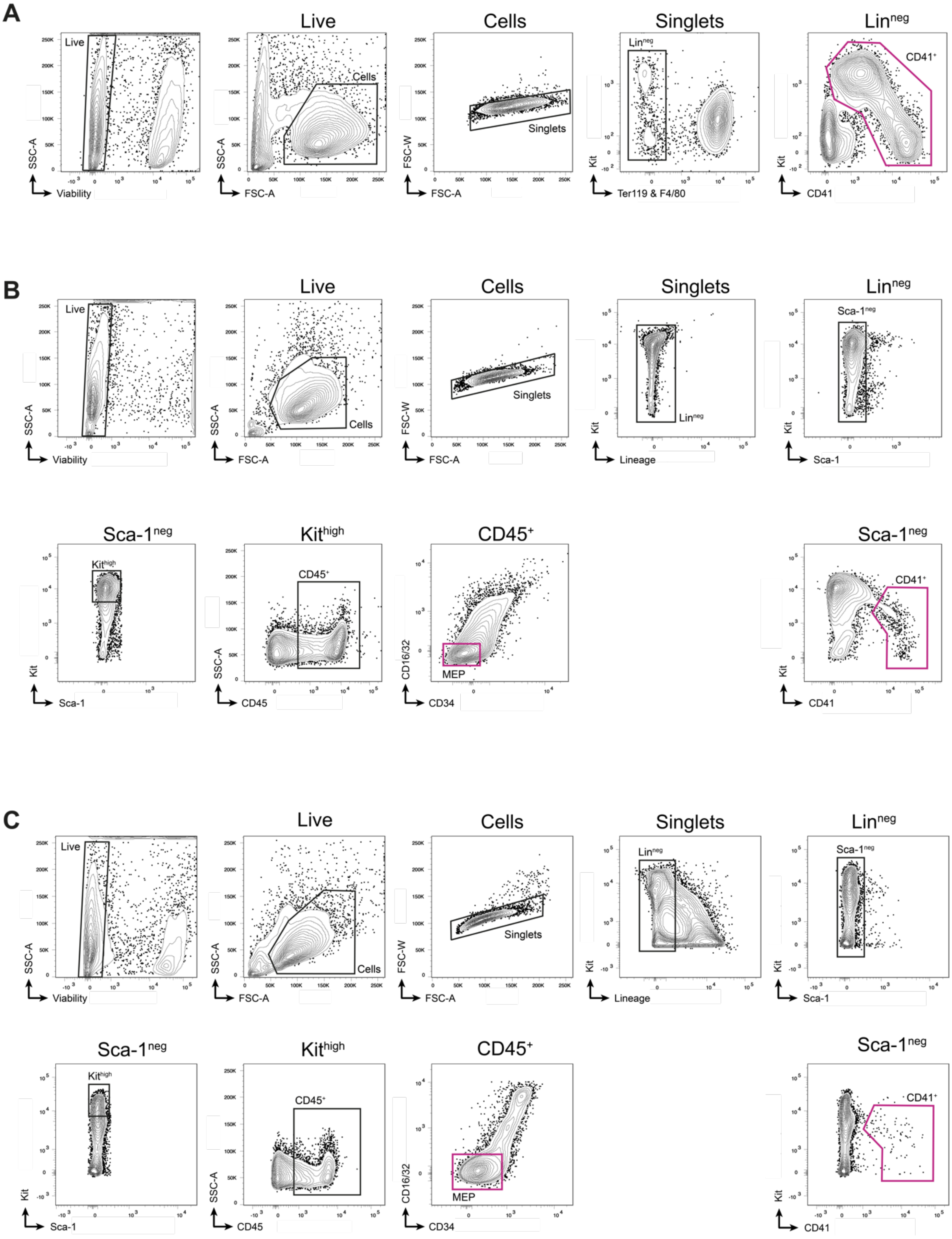


**Supplemental Data Figure 2, to Figure 2.** Gating strategy used for cell sorting prior to single-cell RNA and single-cell ATAC (multiome) sequencing. Purple gates indicate sorted cells from E10.5 yolk sacs (**A**) E12.5 (**B**), and E16.5 (**C**) fetal livers.

#### Supplemental Data Figure 3

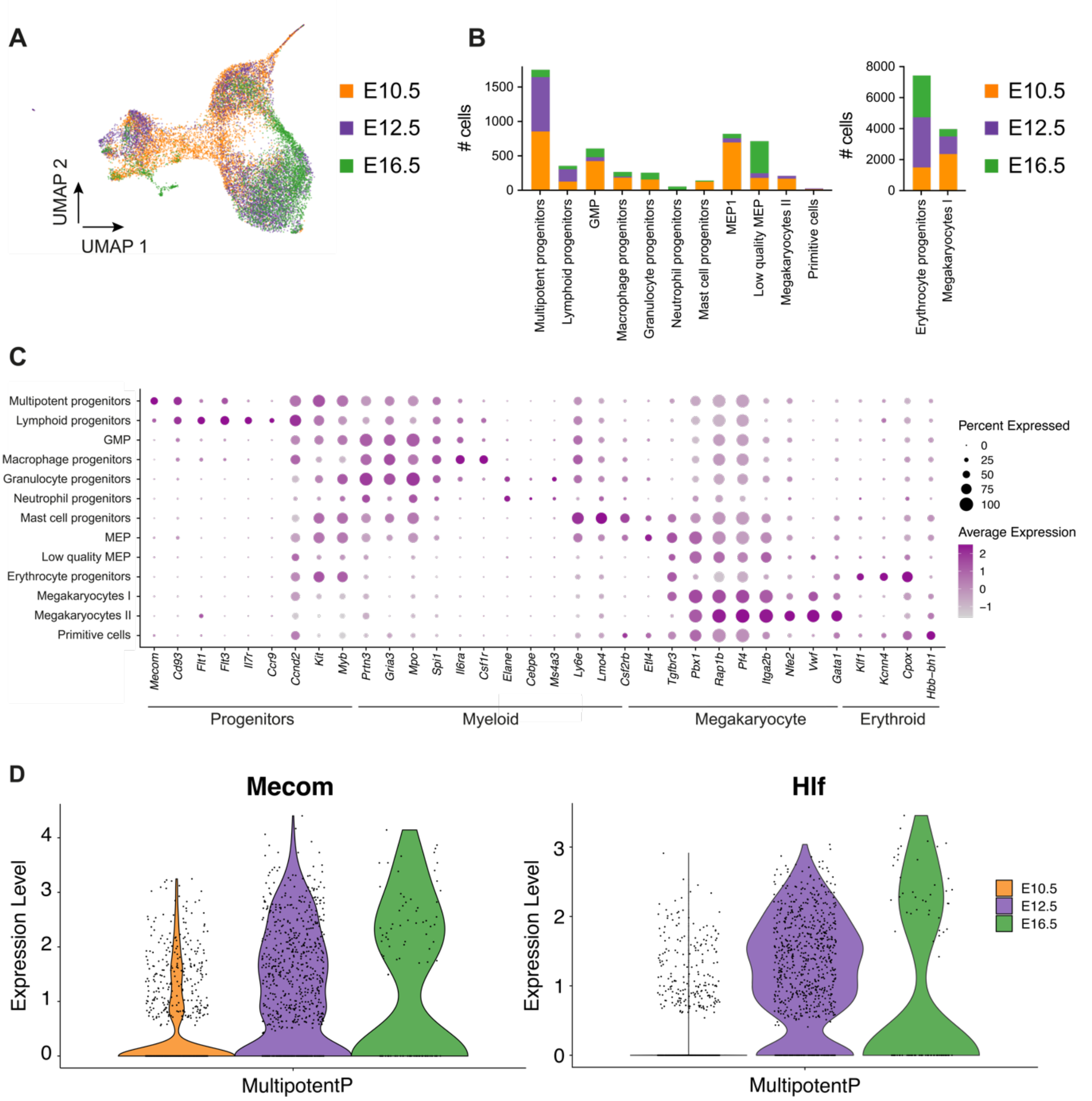


**Supplemental Data Figure 3, to Figure 2.** Integration of single-cell RNA seq data of wild-type (wt) and *Gata1*$^{mCherry}$ (mut) cells from E10.5 yolk sacs, E12.5 fetal livers, and E16.5 fetal livers. (**A, B**) UMAP (**A**) and number of cells per cluster and sample (**B**). E10.5 wt (orange), E12.5 wt (purple), E16.5 wt (green). (**C**) Dot plot showing average expression level (color intensity) and frequency of expression (dot size) of cell type-specific genes among clusters. (**D**) Violin plot illustrating expression levels of *Mecom* (top) and *Hlf* (bottom) in multipotent progenitors (MultipotentP or MPP) at E10.5, E12.5, and E16.5.

**Supplemental Data Figure 4**

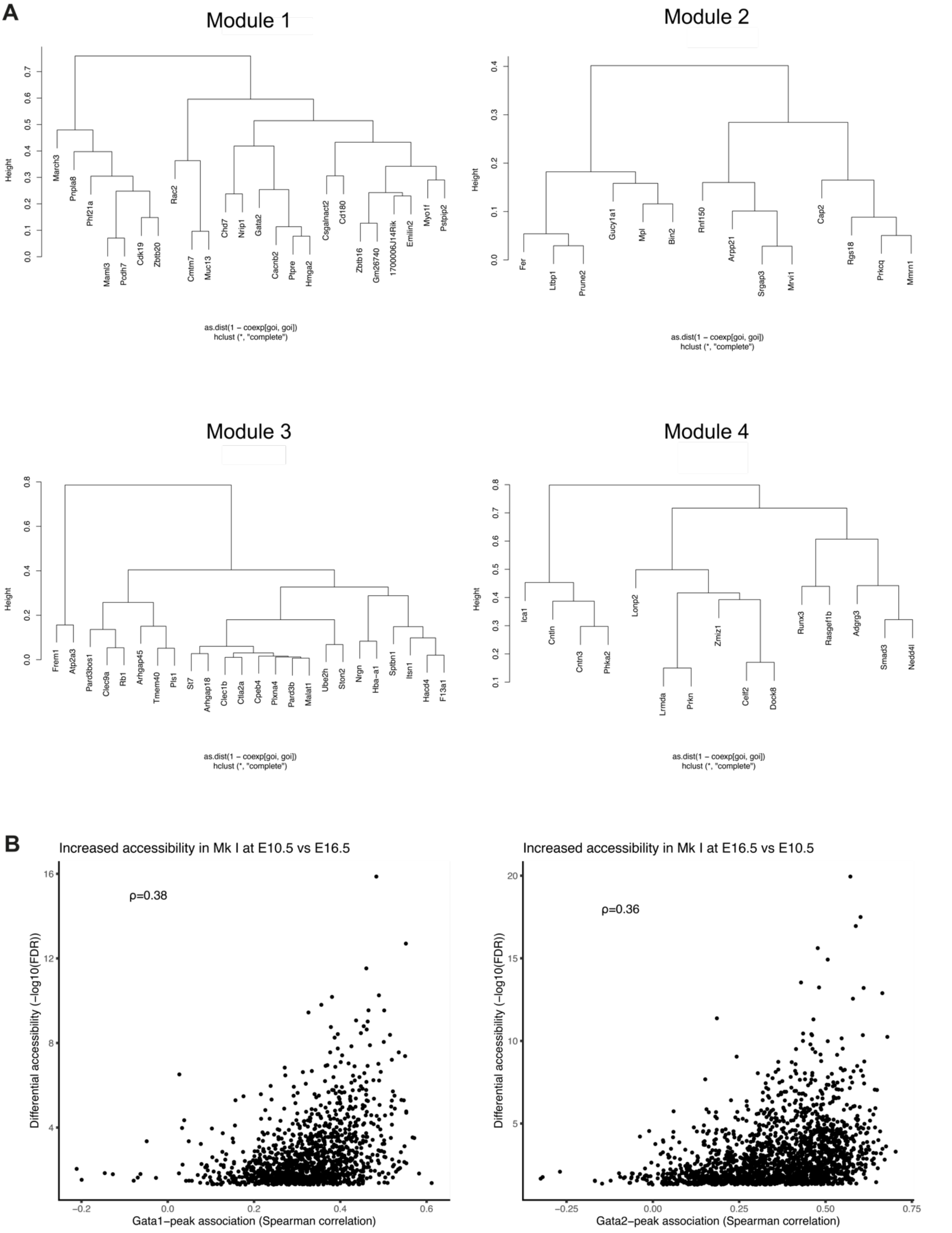


**Supplemental Data Figure 4, to Figure 2.** (**A**) Dendrograms showing top genes among co-expression modules 1-4. (**B**) Plot representing the likelihood that GATA1 (left) or GATA2 (right) is responsible for the opening of the differentially accessible regions at E10.5 (left) or E16.5 (right) compared to E16.5 or E10.5, respectively. <u>x-</u>

axis: spearman correlation of the TF-peak association representing the likelihood that TF is responsible for the opening of the specific chromatin region. y-axis: - log10(FDR(false discovery rate)) of the regions that are differentially accessible at E10.5 or E16.5, compared to E16.5 or E10.5, respectively.

**Supplemental Data Figure 5**

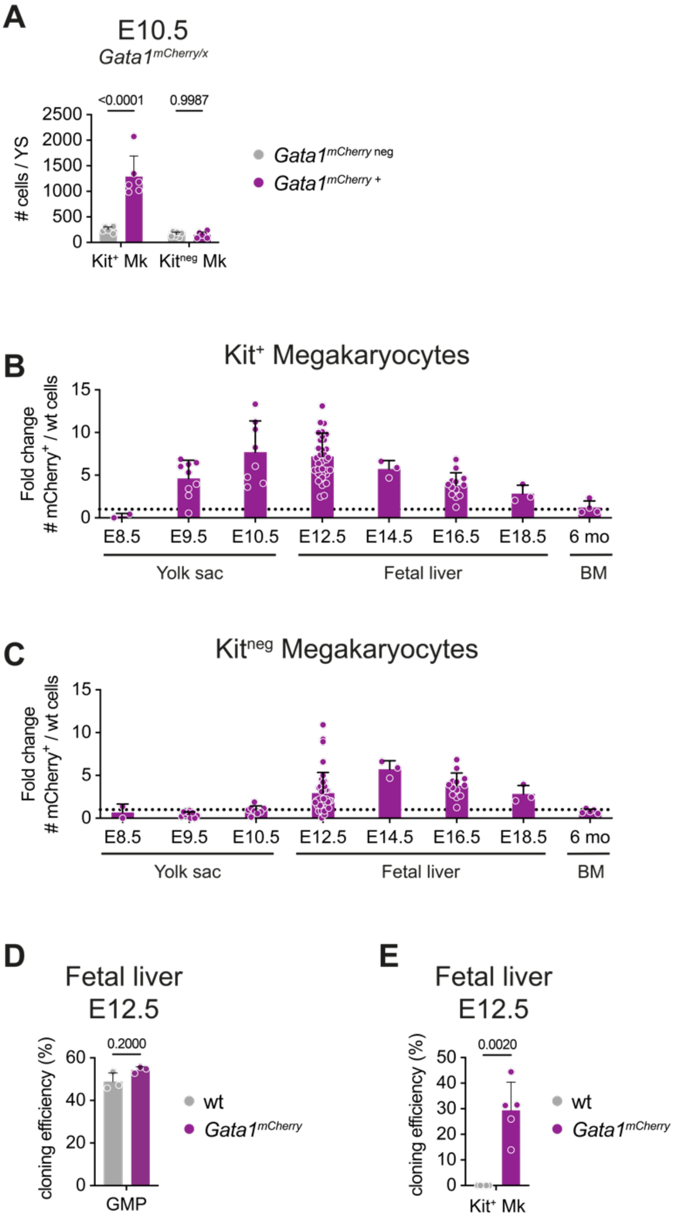


**Supplemental Data Figure 5, to Figure 3.** (**A**) Bar plot showing the number of *Gata1*$^{mCherry\ neg}$ (grey) and *Gata1*$^{mCherry+}$ (purple) Kit$^{+}$ and Kit$^{neg}$ Mks in E10.5 yolk sacs from *Gata1*$^{mCherry/x}$ female embryos. ≥ 3 embryos from ≥ 3 litters per condition. One-way ANOVA with Tukey's multiple comparisons (**B**) Bar plot showing the fold change of numbers of *Gata1*$^{mCherry}$ over wild-type Kit$^{+}$ Mks in the yolk sac (E8.5-E10.5), fetal liver (E12.5-E18.5) and bone marrow (6 months). Dotted line indicates the value "1". ≥ 3 embryos from ≥ 3 litters per condition (**C**) Bar plot showing the fold change of numbers of *Gata1*$^{mCherry}$ over wild-type Kit$^{neg}$ Mks in the yolk sac (E8.5-E10.5), fetal liver (E12.5-E18.5) and bone marrow (6 months). The dotted line indicates the value "1". ≥ 3 embryos from ≥ 3 litters per condition. (**D**) Bar plot showing the cloning efficiency of single immunophenotypic granulocyte-monocyte progenitors (GMPs, $Lin^{neg}$ $Kit^{high}$ $Sca\text{-}1^{neg}$ $CD16/32^{high}$ $CD34^{+}$) in liquid cultures sorted from E12.5 fetal livers from wild-type (grey) or *Gata1*$^{mCherry}$ (purple) embryos. 3 independent litters. Two-tailed Mann-Whitney test. (**E**) Bar plot showing the cloning efficiency of single $Kit^{+}$ Mks sorted from E12.5 fetal livers from wild-type (grey) or *Gata1*$^{mCherry}$ (purple) embryos. 5 independent litters. Mix of liquid and semi-solid cultures Paired one-tailed t-test. Data are represented as mean ± SD.

#### Supplemental Data Figure 6

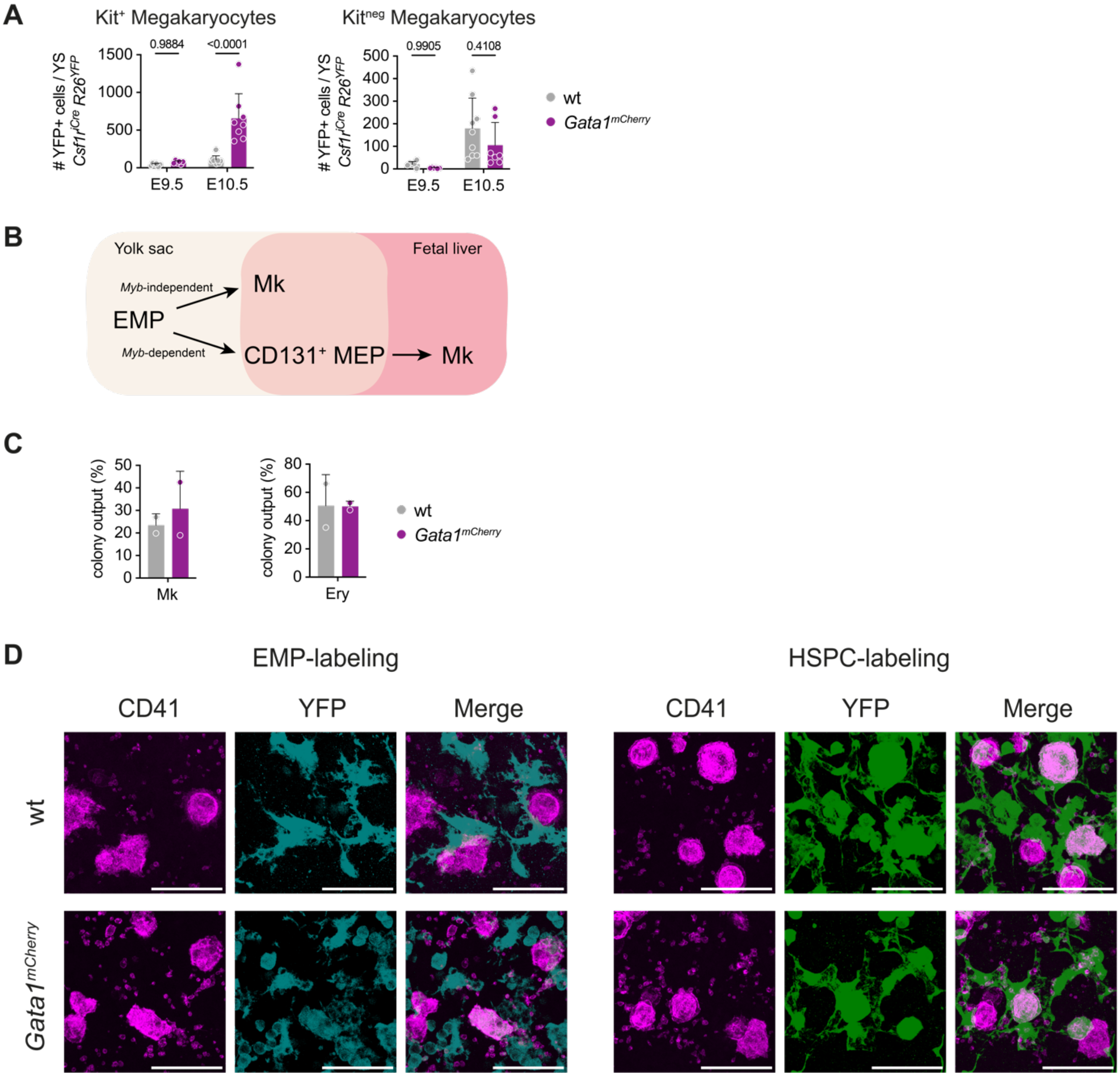


**Supplemental Data Figure 6, to Figure 4.** (**A**) Number of YFP$^+$ Kit$^+$ (left) and YFP$^+$ Kit$^{neg}$ (right) Mks in E9.5 and E10.5 yolk sacs from *Csf1r$^{iCre}$ R26$^{YFP}$ Gata1$^{wild\text{-}type}$* (grey) or *Gata1$^{mCherry}$* (purple) embryos. > 3 embryos from ≥ 3 litters per condition. Two-way ANOVA with Tukey's multiple comparisons: Embryonic stage P-value <0.0001 (left), = 0.0013 (right); Genotype P-value = 0.0002 (left), = 0.2193 (right); Interaction: P-value = 0.0006 (left), 0.4255 (right). Data are represented as mean ± SD. (**B**) Scheme illustrating EMP-derived *Myb*-independent and *Myb*-dependent megakaryocyte differentiation in the yolk sac and fetal liver. (**C**) Bar plot showing megakaryocyte (Mk) and erythrocyte (Ery) output in colonies grown after from Kit$^{high}$, CD41$^+$ CD131$^+$ cells from E10.5 yolk sacs. 2 independent experiments. Data are represented as mean ± SD. (**D**) Immunofluorescence of 42 µm-thick maximum projection of E16.5 fetal livers from wild-type (wt, top) and *Gata1*$^{mCherry}$ (bottom) embryos. Mks are stained with anti-CD41 PE (magenta). EMP-derived cells were fate-mapped in *Csf1r*$^{MeriCreMer}$ *R26*$^{YFP}$ embryos pulsed at E8.5 with 4-OHT (cyan, left). HSPC-derived cells were fate-mapped in *Cdh5*$^{CreERT2}$ *R26*$^{YFP}$ embryos pulsed at E10.5 with 4-OHT (green, right). Scale bar represents 50 µm.

#### Supplemental Data Figure 7

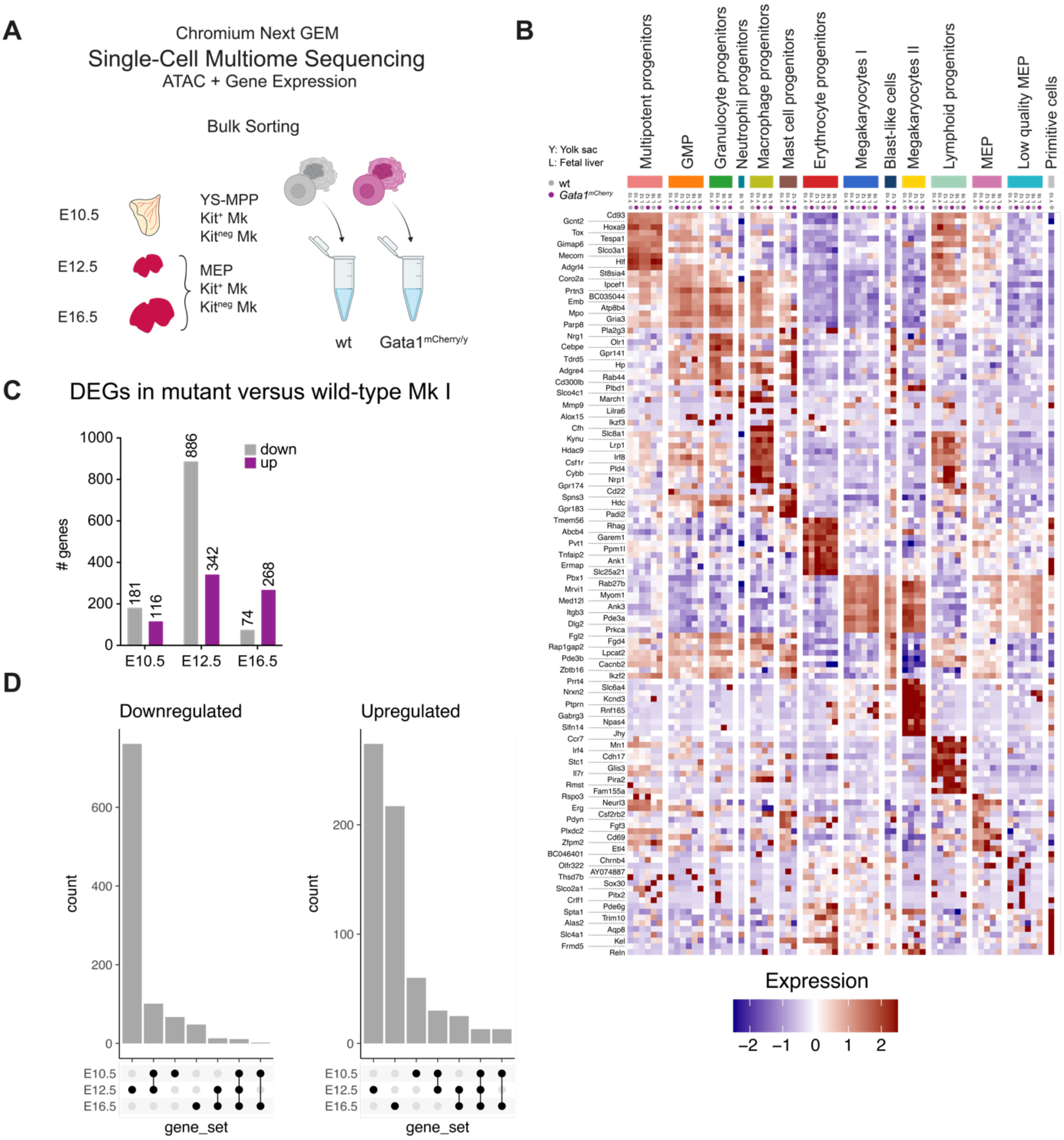


**Supplemental Data Figure 7, to Figure 5.** Integration of single-cell RNA seq data of wild-type (wt) and *Gata1*$^{mCherry}$ (mut) cells from E10.5 yolk sacs, E12.5 fetal livers, and E16.5 fetal livers. (**A**) Scheme illustrating sorting and sequencing strategy. Multipotent progenitors (Ter119$^{neg}$ F4/80$^{neg}$ Kit$^{high}$ CD41$^{int}$), Kit$^{+}$, and Kit$^{neg}$ Mks were sorted from E10.5 yolk sacs. Megakaryocyte-erythrocyte progenitors (MEP), Kit$^{+}$, and Kit$^{neg}$ Mks were sorted from E12.5 and E16.5 fetal livers. Wild-type (male and female) and *Gata1*$^{mCherry}$ (only male) cells were sorted separately at each time point. Cells were lysed, and gene expression and ATAC libraries were prepared from nuclei following the Chromium Next Gem Single-Cell Multiome Sequencing pipeline. (**B**) Heat map showing Top 10 expressed genes per cluster GMP = granulocyte-monocyte progenitor, MEP = megakaryocyte-erythrocyte progenitor. (**C**) Bar plot quantifying the number of up- (purple) and down= (grey) regulated genes between wild-type and *Gata1*$^{mCherry}$ Mks (Mk I) at E10.5, E12.5, and E16.5. (**D**) Up-set plot highlighting down- and upregulated genes in *Gata1*$^{mcherry}$ cells compared to wt cells shared by E10.5, E12.5, and E16.5 samples.

#### Supplemental Data Figure 8

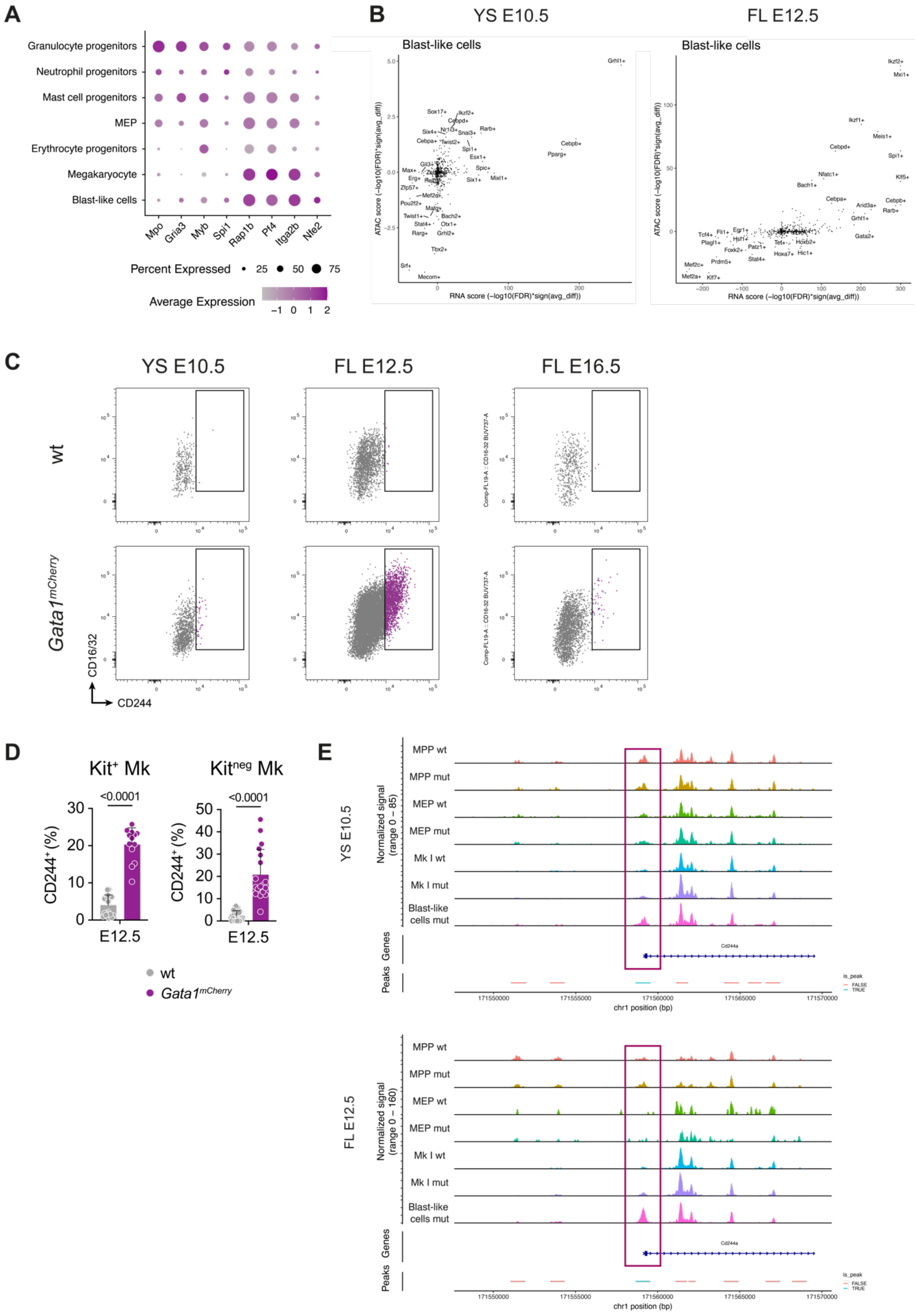


**Supplemental Data Figure 8, to Figure 6.** (**A**) Dot plot showing average expression level (color intensity) and frequency of expression (dot size) of selected genes among clusters. (**B**) RNA- and ATAC scores calculated in Pando analysis indicating transcription factors with increased activity in blast-like cells compared to other clusters in E10.5 yolk sacs and E12.5 fetal livers. (**C**) Dot plots showing gating of $CD244^+$ cells (purple) among $Kit^+$ Mks in E10.5 yolk sacs, E12.5 and E16.5 fetal livers. (**D**) Bar plot showing the frequency of $CD244^+$ cells among $Kit^+$ (left) and $Kit^{neg}$ (right) Mks in E12.5 fetal livers. > 3 embryos from ≥ 3 litters per condition. One-sided Mann-Whitney test. Data are represented as mean ± SD. (**E**) Coverage plot illustrating chromatin accessibility of the *Cd244a* gene in multipotent progenitors (MPP), megakaryocyte-erythrocyte progenitors (MEP), megakaryocytes (Mk I), and blast-like cells from in wild-type (wt) and $\textit{Gata1}^{mCherry}$ (mut) E10.5 yolk sacs and E12.5 fetal livers.

#### Supplemental Data Figure 9

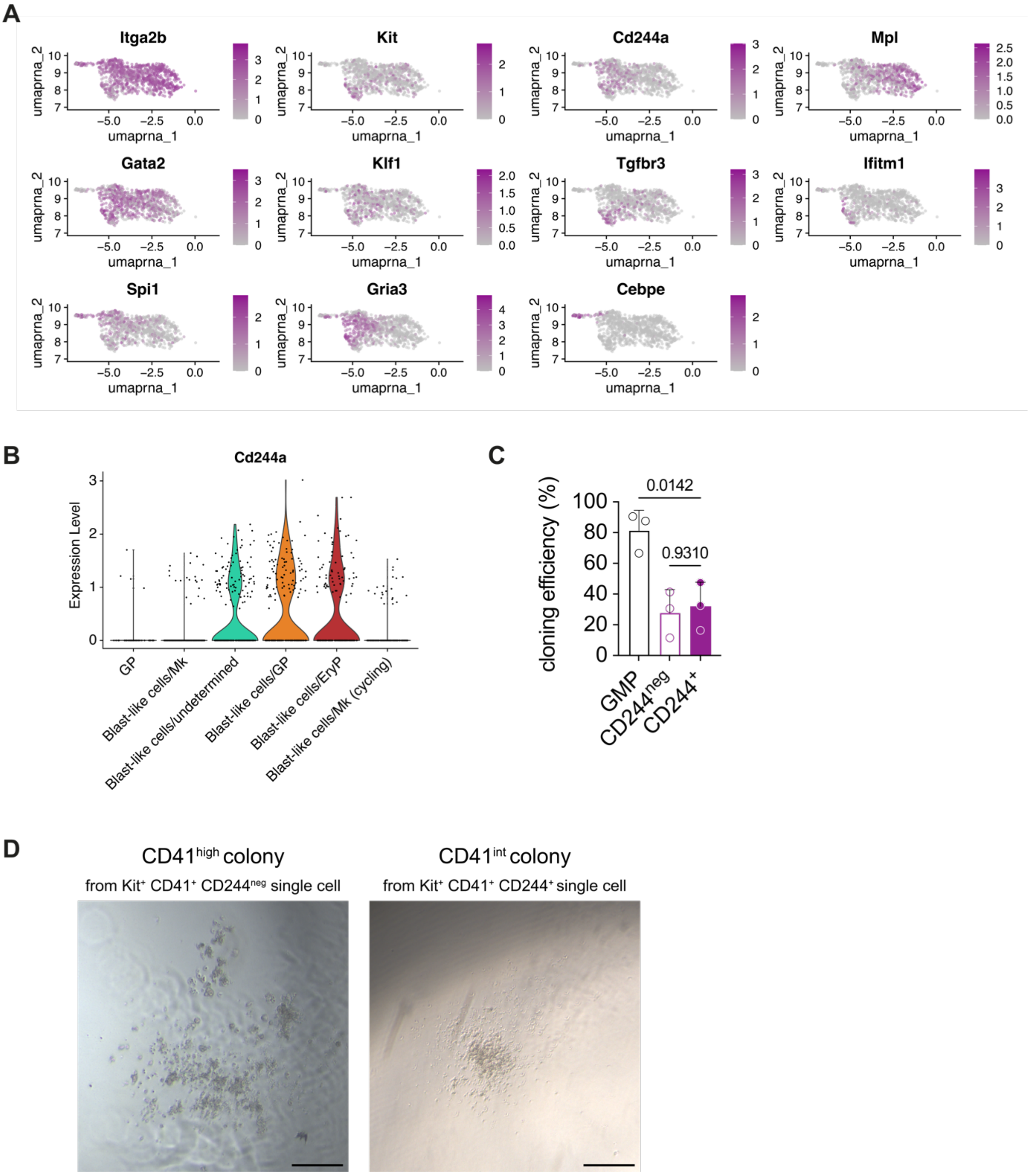


**Supplemental Data Figure 9, to Figure 6.** (**A**) UMAPs displaying gene expression levels in blast-like cells sequenced from E12.5 fetal livers. (**B**) Violin plot showing *Cd244a* expression among sub-clusters in blast-like cells. (**C**) Bar plot showing the cloning efficiency of single immunophenotypic granulocyte-monocyte progenitors (GMPs, $Lin^{neg}$ $Kit^{high}$ $Sca\text{-}1^{neg}$ $CD16/32^{high}$ $CD34^{+}$, black border), $Kit^{+}$ $CD41^{+}$ $CD244^{neg}$ (purple border) or $Kit^{+}$ $CD41^{+}$ $CD244^{+}$ (purple filled) sorted from E12.5 *Gata1*$^{mCherry}$ fetal livers embryos. 3 independent litters. One-way ANOVA with Tukey's multiple comparisons. Data are represented as mean ± SD. (**D**) Bright-field image of CD41$^{high}$ colony grown in methocult for 10 days from a single Kit$^{+}$ CD41$^{+}$ CD244$^{neg}$ cell (left) and CD41$^{int}$ colony grown from a single Kit$^{+}$ CD41$^{+}$ CD244$^{+}$ cell (right) from a E12.5 *Gata1*$^{mCherry}$ fetal liver. Scale bar represents 250 µm.

#### Module 1+2

**Supplemental Table 1.** ChEA 2022 transcription factors predicted to be regulating module 1+2 genes.

| Term | P-value | Adjusted P-value | Odds Ratio | Combined Score | Genes |
|---|---|---|---|---|---|
| GATA1 19941827 ChIP-Seq MEL Mouse | 2,42E-08 | 1,52E-05 | 8,40 | 147,34 | RGS18; ARPP21; PCDH7; ZBTB16; CD180; GATA2; LTBP1; PHF21A; PTPRE; PSTPIP2; NRIP1; RAC2; PRKCQ; SRGAP3; MAML3 |
| GATA1 21571218 ChIP-Seq MEGAKARYOCYTES Human | 5,89E-07 | 0,000186 | 6,13 | 87,98 | RGS18; PCDH7; ZBTB16; PRUNE2; CD180; MPL; ZBTB20; GATA2; LTBP1; PHF21A; CACNB2; PTPRE; FER; MUC13; PRKCQ; MAML3 |
| ISL1 27105846 Chip-Seq CPCs Mouse | 2,84E-06 | 0,000597 | 6,73 | 85,97 | CMTM7; PTPRE; PSTPIP2; PCDH7; ZBTB16; CHD7; NRIP1; CD180; HMGA2; PRKCQ; MAML3; CAP2 |
| WT1 25993318 ChIP-Seq PODOCYTE Human | 3,5E-05 | 0,005529 | 4,29 | 44,02 | CMTM7; ARPP21; PRUNE2; ZBTB20; HMGA2; LTBP1; PHF21A; FER; NRIP1; RAC2; EMILIN2; PRKCQ; SRGAP3; RNF150; MAML3; CAP2 |
| RUNX1 30185409 ChIP-Seq HPC Mouse BoneMarrow Leukemia | 6,91E-05 | 0,006518 | 4,15 | 39,79 | CDK19; PCDH7; ZBTB16; CSGALNACT2; CD180; ZBTB20; GATA2; PHF21A; FER; BIN2; PSTPIP2; NRIP1; MUC13; EMILIN2; MAML3 |
| TAL1 20566737 ChIP-Seq PRIMARY FETAL LIVER ERYTHROID Mouse | 7,02E-05 | 0,006518 | 5,08 | 48,62 | PTPRE; PSTPIP2; ZBTB16; NRIP1; RAC2; MUC13; EMILIN2; HMGA2; PRKCQ; GATA2; LTBP1 |
| NR3C1 27076634 ChIP-Seq BEAS2B Human Lung Inflammation | 7,27E-05 | 0,006518 | 4,00 | 38,16 | CDK19; PCDH7; ZBTB16; CHD7; PRUNE2; CD180; ZBTB20; HMGA2; GATA2; LTBP1; CACNB2; PTPRE; FER; PSTPIP2; NRIP1; RNF150 |
| GATA1 19941827 ChIP-Seq MEL86 Mouse | 8,26E-05 | 0,006518 | 4,99 | 46,87 | RGS18; CDK19; MMRN1; PSTPIP2; ZBTB16; RAC2; PRKCQ; SRGAP3; MAML3; GATA2; PHF21A |
| SOX2 21211035 ChIP-Seq LN229 Gbm | 0,000232 | 0,014864 | 3,80 | 31,77 | RGS18; PNPLA8; PCDH7; CD180; ZBTB20; LTBP1; CACNB2; FER; MMRN1; PSTPIP2; RAC2; SRGAP3; MAML3; CAP2 |
| GATA1 26923725 Chip-Seq HPCs Mouse | 0,000236 | 0,014864 | 14,94 | 124,77 | CDK19; RNF150; PHF21A; MYO1F |
| PAX3-FKHR20663909 ChIP-Seq RHABDOMYOSARCOMA Human | 0,000274 | 0,015696 | 5,58 | 45,80 | CACNB2; PCDH7; CHD7; PRUNE2; ZBTB20; HMGA2; RNF150; MAML3 |
| LMO2 26923725 Chip-Seq HEMOGENIC-ENDOTHELIUM Mouse | 0,000404 | 0,015924 | 4,37 | 34,18 | CDK19; PRUNE2; NRIP1; CD180; PRKCQ; SRGAP3; RNF150; LTBP1; PHF21A; MYO1F |
| MECOM 23826213 ChIP-Seq KASUMI Mouse | 0,000426 | 0,015924 | 4,34 | 33,70 | RGS18; CDK19; PCDH7; CSGALNACT2; CD180; MPL; RAC2; PRKCQ; PHF21A; MYO1F |
| SUZ12 18692474 ChIP-Seq MESCs Mouse | 0,000435 | 0,015924 | 4,33 | 33,52 | CACNB2;PTPRE;ARPP21;PCDH7;ZBTB16;EMILIN2;HMGA2;PRKCQ;RNF150;GATA2 |
| TAL1 30185409 ChIP-Seq HPC Mouse BoneMarrow Leukemia | 0,000438 | 0,015924 | 3,84 | 29,69 | RGS18; PTPRE; FER; BIN2; PCDH7; ZBTB16; PRUNE2; NRIP1; CSGALNACT2; ZBTB20; EMILIN2; PHF21A |
| AR 22383394 ChIP-Seq PROSTATE CANCER Human | 0,000456 | 0,015924 | 4,30 | 33,10 | RGS18; CDK19; FER; PCDH7; NRIP1; CD180; ZBTB20; SRGAP3; MAML3; CAP2 |
| SMAD4 21799915 ChIP-Seq A2780 Human | 0,000463 | 0,015924 | 3,81 | 29,28 | RGS18; CACNB2; BIN2; PCDH7; CHD7; NRIP1; RAC2; ZBTB20; HMGA2; SRGAP3; LTBP1; PHF21A |

**Supplemental Table 2.** GO Biological Processes (2023) predicted to be modulated by module 1+2 genes.

| Term | P-value | Adjusted P-value | Odds Ratio | Combined Score | Genes |
|---|---|---|---|---|---|
| Regulation Of Platelet Aggregation (GO:0090330) | 1,41E-05 | 0,006 | 77,688 | 867,512 | MMRN1; EMILIN2; PRKCQ |
| Positive Regulation Of Platelet Aggregation (GO:1901731) | 1,91E-04 | 0,038 | 123,167 | 1054,544 | MMRN1; EMILIN2 |
| Positive Regulation Of Homotypic Cell-Cell Adhesion (GO:0034112) | 4,15E-04 | 0,055 | 79,159 | 616,469 | MMRN1; EMILIN2 |
| Fc-epsilon Receptor Signaling Pathway (GO:0038095) | 7,93E-04 | 0,060 | 55,394 | 395,517 | FER; PRKCQ |
| Positive Regulation Of Vasculature Development (GO:1904018) | 1,01E-03 | 0,060 | 16,855 | 116,272 | EMILIN2; HMGA2; GATA2 |
| Negative Regulation Of Insulin Receptor Signaling Pathway (GO:0046627) | 1,11E-03 | 0,060 | 46,153 | 313,996 | PTPRE; PRKCQ |
| Fc Receptor Signaling Pathway (GO:0038093) | 1,11E-03 | 0,060 | 46,153 | 313,996 | FER; PRKCQ |
| Negative Regulation Of Cellular Response To Insulin Stimulus (GO:1900077) | 1,20E-03 | 0,060 | 44,304 | 298,064 | PTPRE; PRKCQ |
| Positive Regulation Of Angiogenesis (GO:0045766) | 1,49E-03 | 0,066 | 14,665 | 95,467 | EMILIN2; HMGA2; GATA2 |
| Regulation Of Lamellipodium Assembly (GO:0010591) | 1,79E-03 | 0,071 | 35,719 | 225,974 | FER; RAC2 |

#### Module 3+4

**Supplemental Table 3.** ChEA 2022 transcription factors predicted to be regulating module 3+4 genes.

| Term | P-value | Adjusted P-value | Odds Ratio | Combined Score | Genes |
|---|---|---|---|---|---|
| SMAD3 21741376 ChIP-Seq HESCs Human | 1,82E-06 | 0,001177 | 7,667 | 101,329 | SMAD3; RASGEF1B; ZMIZ1; DOCK8; ARHGAP18; NEDD4L; TMEM40; FREM1; PLXNA4; CNTLN; CPEB4 |
| GATA2 19941826 ChIP-Seq K562 Human | 6,66E-06 | 0,002153 | 5,718 | 68,163 | UBE2H; SMAD3; DOCK8; ITSN1; ICA1; STON2; PARD3B; RASGEF1B; ZMIZ1; ST7; SPTBN1; PLS1; CPEB4 |
| WT1 25993318 ChIP-Seq PODOCYTE Human | 1,15E-05 | 0,002452 | 4,561 | 51,876 | UBE2H; SMAD3; CELF2; DOCK8; ITSN1; ICA1; NEDD4L; TMEM40; PHKA2; NRGN; PARD3B; RASGEF1B; ZMIZ1; LONP2; CNTN3; FREM1; PLXNA4 |
| SCL 21571218 ChIP-Seq MEGAKARYOCYTES Human | 1,52E-05 | 0,002452 | 5,607 | 62,222 | SMAD3; RASGEF1B; ZMIZ1; ST7; ATP2A3; TMEM40; STON2; RUNX3; PLXNA4; SPTBN1; NRGN; CPEB4 |
| SMARCA4 23332759 ChIP-Seq OLIGODENDROCYTES Mouse | 2,12E-05 | 0,002743 | 4,827 | 51,947 | CELF2; DOCK8; ICA1; ARHGAP18; NEDD4L; CLEC9A; STON2; PARD3B; ST7; CNTN3; FREM1; PLS1; CNTLN; CPEB4 |
| STAT3 23295773 ChIP-Seq U87 Human | 6,73E-05 | 0,007258 | 4,133 | 39,703 | DOCK8; ARHGAP18; F13A1; NEDD4L; RUNX3; PARD3B; ST7; LONP2; CNTN3; CLEC1B; FREM1; PLXNA4; SPTBN1; PLS1; CNTLN |
| SMAD4 21741376 ChIP-Seq HESCs Human | 0,000147 | 0,013562 | 4,351 | 38,406 | SMAD3; RASGEF1B; ZMIZ1; DOCK8; ARHGAP18; NEDD4L; TMEM40; CNTN3; RUNX3; FREM1; PLXNA4; CPEB4 |
| SMRT 27268052 Chip-Seq Bcells Human | 0,000191 | 0,015471 | 4,478 | 38,338 | RB1; UBE2H; CELF2; ARHGAP18; F13A1; NEDD4L; TMEM40; RUNX3; PLXNA4; SPTBN1; CPEB4 |
| GATA1 30185409 ChIP-Seq HPC Mouse BoneMarrow Leukemia | 0,000237 | 0,016855 | 5,690 | 47,498 | UBE2H; ZMIZ1; ITSN1; ATP2A3; LONP2; STON2; RUNX3; CPEB4 |
| OLIG2 23332759 ChIP-Seq OLIGODENDROCYTES Mouse | 0,000261 | 0,016855 | 4,308 | 35,557 | CELF2; ST7; ARHGAP18; NEDD4L; LONP2; CNTN3; FREM1; PLS1; CNTLN; PARD3B; CPEB4 |
| HTT 18923047 ChIP-ChIP STHdh Human | 0,000296 | 0,017405 | 9,670 | 78,577 | ARHGAP18; F13A1; TMEM40; CLEC9A; CPEB4 |
| Nerf2 26677805 Chip-Seq MACROPHAGESS Mouse | 0,000559 | 0,030116 | 4,972 | 37,238 | SMAD3; RASGEF1B; DOCK8; ATP2A3; NEDD4L; CLEC9A; STON2; CPEB4 |
| ZNF217 24962896 ChIP-Seq MCF-7 Human | 0,000679 | 0,03208 | 4,378 | 31,941 | UBE2H; SMAD3; ZMIZ1; ICA1; STON2; PLXNA4; SPTBN1; PARD3B; CPEB4 |
| ISL1 27105846 Chip-Seq CPCs Mouse | 0,000694 | 0,03208 | 4,364 | 31,737 | ZMIZ1; CELF2; ATP2A3; ARHGAP18; STON2; RUNX3; PHKA2; FREM1; PARD3B |
| HOXA2 22223247 ChIP-Seq E11.5 EMBRYO Mouse | 0,000913 | 0,039399 | 51,326 | 359,192 | ARHGAP18; FREM1 |
| PPARG 20176806 ChIP-Seq MACROPHAGES Mouse | 0,000987 | 0,039904 | 4,534 | 31,378 | RASGEF1B; ZMIZ1; CELF2; DOCK8; ITSN1; ARHGAP18; F13A1; FREM1 |
| CEBPB 20513432 ChIP-Seq MACROPHAGES Mouse | 0,00184 | 0,066295 | 3,760 | 23,679 | UBE2H; DOCK8; ITSN1; ST7; ARHGAP18; NEDD4L; CLEC1B; NRGN; CPEB4 |

**Supplemental Table 4.** GO Biological Processes (2023) predicted to be modulated by module 3+4 genes.

| Term | P-value | Adjusted P-value | Odds Ratio | Combined Score | Genes |
|---|---|---|---|---|---|
| Regulation Of Small GTPase Mediated Signal Transduction (GO:0051056) | 7,28E-05 | 0,036 | 20,483 | 195,162 | DOCK8; ITSN1; ARHGAP18; ARHGAP45 |
| Protein K48-linked Ubiquitination (GO:0070936) | 3,17E-04 | 0,065 | 25,452 | 205,037 | PRKN; UBE2H; NEDD4L |
| Positive Regulation Of Protein Localization To Membrane (GO:1905477) | 4,53E-04 | 0,065 | 22,428 | 172,689 | PRKN; SPTBN1; PLS1 |
| Positive Regulation Of Extracellular Matrix Organization (GO:1903055) | 0,001 | 0,065 | 69,257 | 522,697 | RB1; SMAD3 |
| Clathrin-Dependent Endocytosis (GO:0072583) | 0,001 | 0,076 | 44,304 | 298,064 | ITSN1; STON2 |
| Small GTPase Mediated Signal Transduction (GO:0007264) | 0,001 | 0,076 | 15,612 | 104,355 | RB1; RASGEF1B; ARHGAP18 |
| Regulation Of Protein Localization To Membrane (GO:1905475) | 0,001 | 0,076 | 42,598 | 283,480 | PRKN; SPTBN1 |
| Protein K11-linked Ubiquitination (GO:0070979) | 0,001 | 0,076 | 41,019 | 270,085 | PRKN; UBE2H |
| Regulation Of Protein Modification By Small Protein Conjugation Or Removal (GO:1903320) | 0,001 | 0,076 | 41,019 | 270,085 | PRKN; ZMIZ1 |
| Regulation Of Cell Motility (GO:2000145) | 0,002 | 0,081 | 14,173 | 90,922 | ADGRG3; ARHGAP18; PLXNA4 |
